# Nuclear and mitochondrial phylogenomics of the Diplostomoidea and Diplostomida (Digenea, Platyhelminthes)^1^

**DOI:** 10.1101/333518

**Authors:** Sean A. Locke, Alex Van Dam, Monica Caffara, Hudson Alves Pinto, Danimar López-Hernández, Christopher Blanar

**Affiliations:** University of Puerto Rico at Mayagüez, Department of Biology, Box 9000, Mayagüez, Puerto Rico 00681–9000; Department of Veterinary Medical Sciences, Alma Mater Studiorum University of Bologna, Via Tolara di Sopra 50, 40064 Ozzano Emilia (BO), Italy; Departament of Parasitology, Instituto de Ciências Biológicas, Universidade Federal de Minas Gerais, Belo Horizonte, Minas Gerais, Brazil.; Nova Southeastern University, 3301 College Avenue, Fort Lauderdale, Florida, USA 33314-7796.

**Keywords:** Strigeida, Diplostomida, Phylogenomics, Metacercaria, Nuclear-mitochondrial discordance, *Cotylurus*, *Hysteromorpha*, *Alaria*

## Abstract

Higher systematics within the Digenea, Carus 1863 have been relatively stable since a phylogenetic analysis of partial nuclear ribosomal markers (rDNA) led to the erection of the Diplostomida Olson, Cribb, Tkach, Bray, and Littlewood, 2003. However, recent mitochondrial (mt) genome phylogenies suggest this order might be paraphyletic. These analyses show members of two diplostomidan superfamilies are more closely related to the Plagiorchiida La Rue, 1957 than to other members of the Diplostomida. In one of the groups implicated, the Diplostomoidea Poirier, 1886, a recent phylogeny based on mt DNA also indicates the superfamily as a whole is non-monophyletic. To determine if these results were robust to additional taxon sampling, we analyzed mt genomes from seven diplostomoids in three families. To choose between phylogenetic alternatives based on mt genomes and the prior rDNA-based topology, we also analyzed hundreds of ultra-conserved elements (UCEs) assembled from shotgun sequencing. The Diplostomida was paraphyletic in the mt genome phylogeny, but supported in the UCE phylogeny. We speculate this mitonuclear discordance is related to ancient, rapid radiation in the Digenea. Both UCEs and mt genomes support the monophyly of the Diplostomoidea and show congruent relationships within it. The Cyathocotylidae Mühling, 1898 are early diverging descendants of a paraphyletic clade of Diplostomidae Poirier, 1886, in which were nested members of the Strigeidae Railliet, 1919; the results support prior suggestions that the Crassiphialinae Sudarikov, 1960 will rise to the family level. Morphological traits of diplostomoid metacercariae appear to be more useful for differentiating higher taxa than those of adults. We describe a new species of *Cotylurus* Szidat, 1928, resurrect a species of *Hysteromorpha* Lutz, 1931, and find support for a species of *Alaria* Schrank, 1788 of contested validity. Complete rDNA operons are provided as a resource for future studies.

## 1. Introduction

Early efforts to organize the higher taxonomy of digenetic trematodes relied mainly on external morphology of adults (reviewed by La Rue, 1957). As life cycles became better known, Cort (1917), Stunkard (1946) and others increasingly emphasized other characters, particularly cercarial morphology. Using a variety of methods, different authors produced conflicting hypotheses and higher classifications, but many concluded a close relationship exists among members of the Diplostomoidea Poirier, 1886, the Clinostomatoidea Lühe, 1901, and the Schistosomatoidea Stiles and Hassall, 1898 (Brooks et al., 1985; Dubois, 1970a; Gibson, 1996; La Rue, 1957; Pearson, 1972). Most authors assigned these and other superfamilies to the Strigeida (=Strigeatoidea) La Rue 1957, which are characterized by furcocercous cercariae that penetrate hosts (Gibson and Bray, 1994). The close relationship among strigeids, clinostomes and schistosomes was supported by a phylogenetic analysis of partial nuclear ribosomal markers (rDNA) from digeneans in 77 families (Olson et al., 2003). In this analysis, however, several other families in the Strigeida fell within the Plagiorchiida La Rue, 1957, leading Olson et al. (2003) to erect the order Diplostomida, which now includes diplostomoids, clinostomatids, and schistosomatoids.

The work of Olson et al. (2003) created stability in higher systematics within the Digenea. For example, the subsequent studies and future research directions discussed by Kostadinova and Pérez-del-Olmo (2014) are mainly limited to intra-ordinal relationships. However, recent work raises questions about the status of the Diplostomida (Fig. 1). Separate phylogenetic analyses of mt genomes show two diplostomidans (*Clinostomum* Leidy, 1856, *Diplostomum* von Nordmann, 1832) are more closely related to the Plagiorchiida than to other members of the Diplostomida (Brabec et al., 2015; Briscoe et al., 2016; Chen et al., 2016; see also Fig. 5 in Park, 2007). Brabec et al. (2015) argued that data from additional taxa are needed before biological and taxonomic implications can be judged. However, it is well to note that the mt phylogenies were based on alignments of considerably more characters than the rDNA phylogeny of Olson et al. (2003). Moreover, mt genome analyses now collectively include two of three diplostomidan superfamilies, and statistical support for the alliance of non-schistosome diplostomidans with the Plagiorchiida has increased with increased taxon sampling (support of the key nodes of 0.77 in Brabec et al., 2015; 1.0 in Chen et al., 2016; 0.93 in Briscoe et al., 2016). More generally, mtDNA has been useful in revealing ordinal relationships in other Platyhelminthes (Waeschenbach et al., 2012). Further evaluation of this discordance (Fig. 1) was one of two goals of our study.

**Fig 1.**
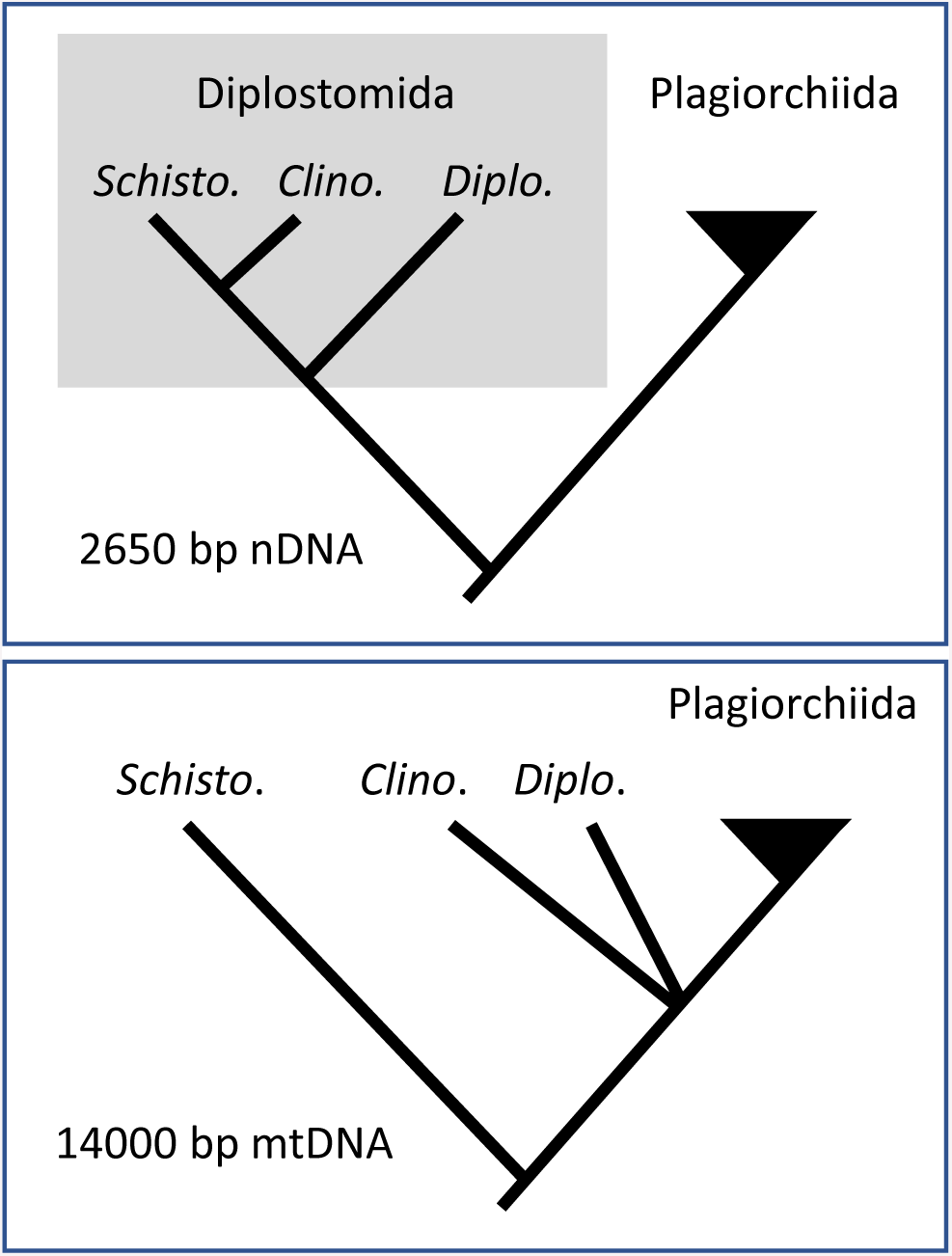
Schematic of phylogenetic conflict in the Digenea emerging from prior studies. Analysis of nuclear rDNA (nDNA) indicates *Diplostomum, Clinostomum*, and *Schistosoma* belong to the Diplostomida (Olson et al., 2003), while mitochondrial genomes (mtDNA) indicate *Diplostomum* and *Clinostomum* belong to the Plagiorchiida (Brabec et al., 2015; Briscoe et al., 2016; Chen et al., 2016). A polytomy occurs in the mitochondrial phylogeny because *Diplostomum* and *Clinostomum* have not been included together in prior analysis.

Another was to estimate relationships among members of one superfamily in the Diplostomida, the Diplostomoidea, which are characterized by the tribocytic organ, a holdfast absent in other digeneans. Six families recognized or erected by Dubois (1938, 1970b) are in wide use, although problems with these classifications are often noted. Shoop’s (1989) cladistic analysis indicated the Diplostomidae Poirier, 1886 was paraphyletic. Both Shoop (1989) and Niewiadomska (2002a) agreed that the morphological type of the metacercaria appears to reflect higher relationships in the Diplostomoidea, but the higher classifications of Dubois (1938, 1970b), based largely on the class of hosts and morphology adults, did not take this into account. Most molecular phylogenetic studies indicate the superfamily is monophyletic, but find the two largest families, the Strigeidae Railliet, 1919 and Diplostomidae, are not (Blasco-Costa and Locke, 2017; Dzikowski et al., 2004; Olson et al., 2003). One analysis of partial cytochrome *c* oxidase 1 (CO1) recovered the Cyathocotylidae Mühling, 1898 (Diplostomoidea) within the Schistosomatoidea (Hernández-Mena et al., 2017), which suggests even the superfamily could be non-monophyletic.

Two obstacles currently prevent molecular assessment of higher taxonomy within the Diplostomoidea, its position within the Digenea, and implications for the order Diplostomida. The first is poor taxonomic and geographic coverage (Blasco-Costa and Locke, 2017). Recent work is encouraging, with studies including members of the Proterodiplostomidae Dubois, 1936 (Hernández-Mena et al., 2017), Brauninidae Wolf, 1903 and Cyathocotylidae (Blasco-Costa and Locke, 2017; Fraija-Fernández et al., 2015), and less well-sampled regions (Blasco-Costa et al., 2016; Chaudhary et al., 2017; López-Hernández et al., 2018; Sereno-Uribe et al., 2018).

Another issue is that accumulating molecular data may not be sufficiently rich in characters to resolve higher relationships. Most analyses of intra-diplostomoid relations are based on sequences of DNA from one or two loci totalling less than 2000 bp in length (e.g., Blasco-Costa and Locke, 2017, and references therein). This seems sufficient for specimen identification, discrimination of species, and species membership in genera (e.g., Chibwana et al. 2013; Locke et al. 2015; López-Hernández et al., 2018), but relationships among genera and families are less resolved. For example, there is inconsistency and often little support in the polarity among *Alaria, Apharyngostrigea,* and *Cardiocephaloides* in fig. 3 in Olson et al. (2003), fig. 1 in Fraija-Fernández et al. (2015), and fig. 2 in Hernández-Mena et al. (2017), which are all phylogenies based on concatenated sequences of rDNA subunits. The deep conflict between recent mt genome phylogenies and those based on shorter nuclear DNA sequences (Fig. 1) may also be related to differences in the number of informative characters.

**Fig 2.**
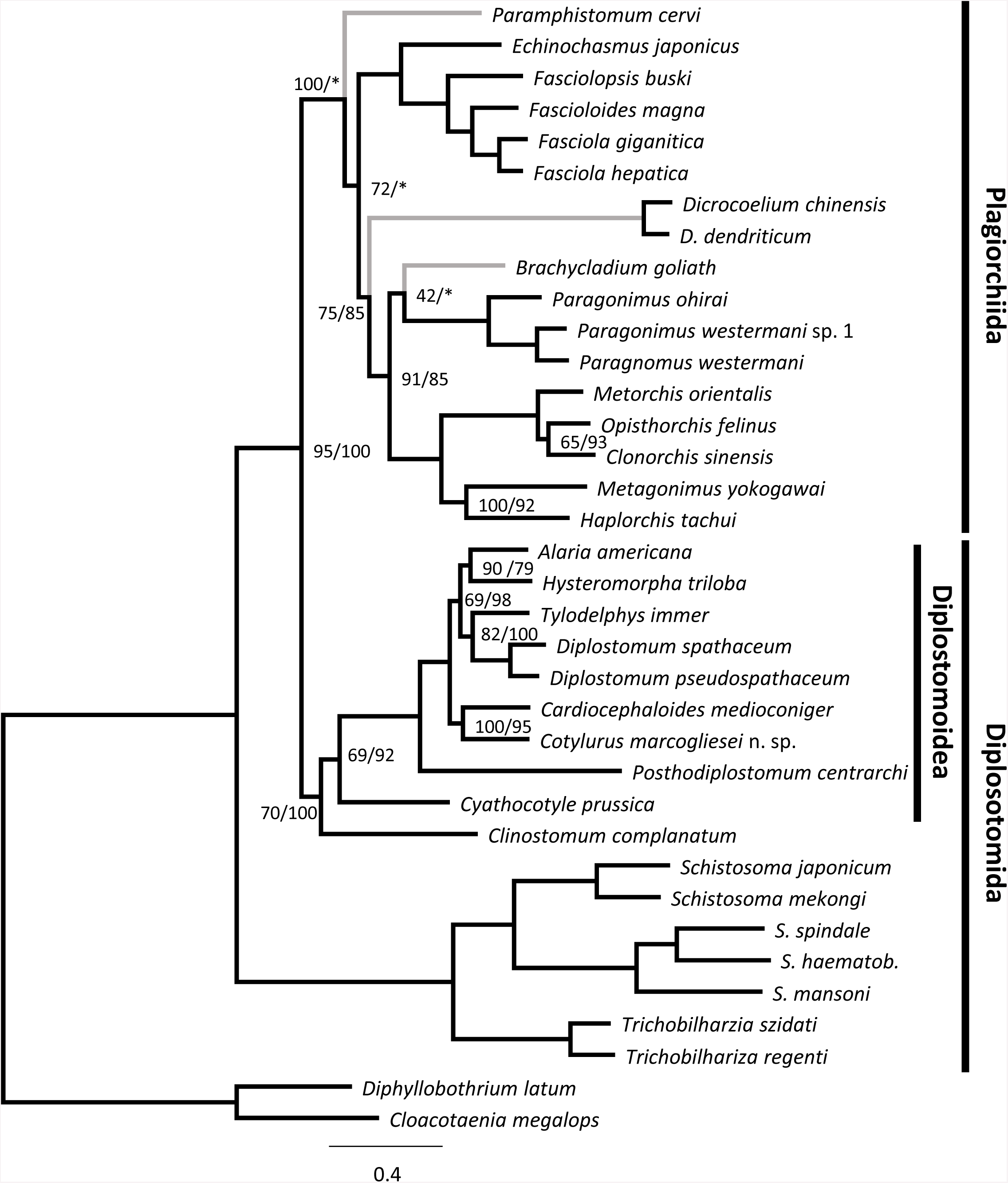
Phylogenetic analysis of seven representatives of the Diplostomoidea and 29 other members of the Platyhelminthes, estimated using maximum likelihood based on 5647 variable sites in 13 protein-coding genes in the mitochondrion. Nodes are labelled with support from 1000 bootstrap replicates, with support from analysis of translated amino acids (supplementary Fig. 2) after the slash. Unlabelled nodes indicate support of 100/100. An asterisk and grey branch indicate topological inconsistency with analysis of amino acids. GenBank accessions of sequences from other studies are AF215860.1, AF216697.1, AF217449.1, AF219379.2, DQ157222.2, DQ157223.1, DQ859919.1, DQ985706.1, EU921260.2, FJ381664.2, HE601612.1, KC330755.1, KF214770.1, KF318786.1, KF318787.1, KF475773.1, KF543342.1, KM280646.1, KM923964.1, KP844722.1, KR269763.1, KR269764.1, KR703278.1, KT239342.1, KU060148.1, KU641017.1, KX169163.1, KX765277.1, MF136777.1.

**Fig 3.**
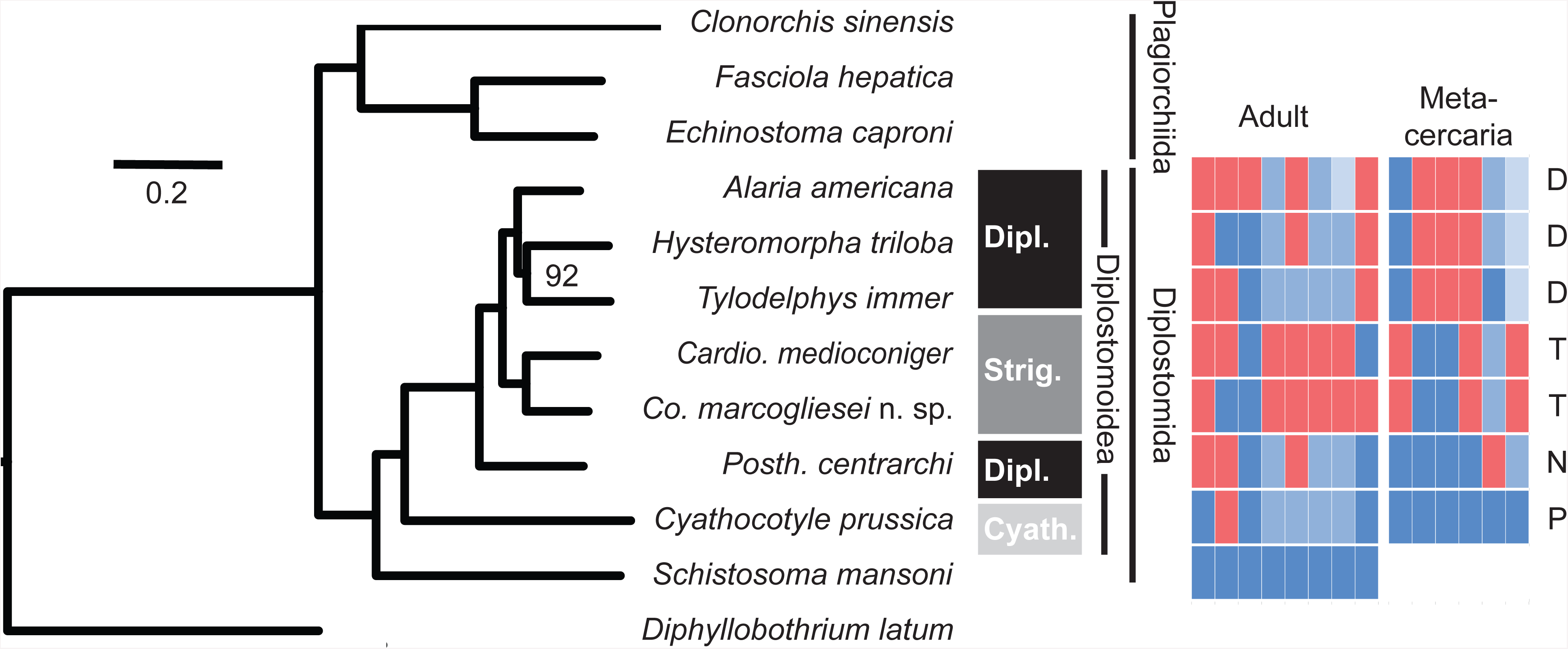
Maximum likelihood analysis of 517 conserved genetic elements from seven members of the Diplostomoidea, and other Platyhelminthes. The gray or black shaded boxes indicate families recognized by Dubois (Dubois, 1938, 1970b) and Niewiadomska (Niewiadomska, 2002a). Colors in matrix at right encode morphological and life history characters separated by developmental stage (supplementary Table 1). Metacercarial morphotypes indicated at far right are: P=prohemistomulum, D=diplostomulum, N=neascus, T=tetracotyle. The alignment is 234,783 bp in length, with 149,881 (64%) invariant and 84,902 (36%) variable sites. The analysis was based on 90% matrix occupancy. Nodes had 100% support in 1000 bootstrap replicates, except where indicated. Accessions of non-diplostomoid genomes obtained from Wormbase (Howe et al., 2016) are PRJEB1206, PRJDA72781, PRJEB6687, PRJEB1207.

Here we attempt to progress on both fronts. We increase the number and diversity of mitochondrial genomes from the Diplostomoidea, in order to explore discordant results of recent phylogenetic studies, namely the possible placement of the Cyathocotylidae outside the Diplostomoidea (Hernández-Mena et al., 2017), and the Diplostomidae as a basal lineage in the order Plagiorchiida (Brabec et al., 2015). Both results originate from analysis of mtDNA, and we obtained data to determine if these patterns were robust to additional taxon sampling. To decide between what were likely to be conflicting topologies based on mtDNA and rDNA, we employed phylogenomic analyses of ultra-conserved elements (UCEs).

Although we set out to work on higher relationships among a small number of representative specimens, we were aware that the diversity and identity of diplostomoid species often differs from initial suspicions based on morphology (Blasco-Costa and Locke, 2017). We therefore supported our identifications with morphological comparisons and analysis of additional DNA sequences from closely related species, which led to several findings related to the alpha taxonomy of the seven specimens of principal interest.

## 2. Materials and Methods

### 2.1 Specimen collection and identification

Twenty-five specimens in good condition were selected for potential Illumina shotgun sequencing (described below). These worms had been stored in ethanol and identified to genus, and were chosen to represent major clades in Blasco-Costa and Locke (2017). DNA was extracted from individual worms, or subsamples, using Qiagen’s DNEasy blood and tissue kit (GmbH, Germany), following the manufacturer’s protocol with two 200-μL elutions. Of these 25 worms, we selected seven (Table 1) after measuring DNA concentration with Nano-drop (0.5-22 ng/μl in 400 μl) and Qubit (0-4.96 ng/μl) and excluding samples with evidence of DNA degradation (Bioanalyzer).

**Table 1:**
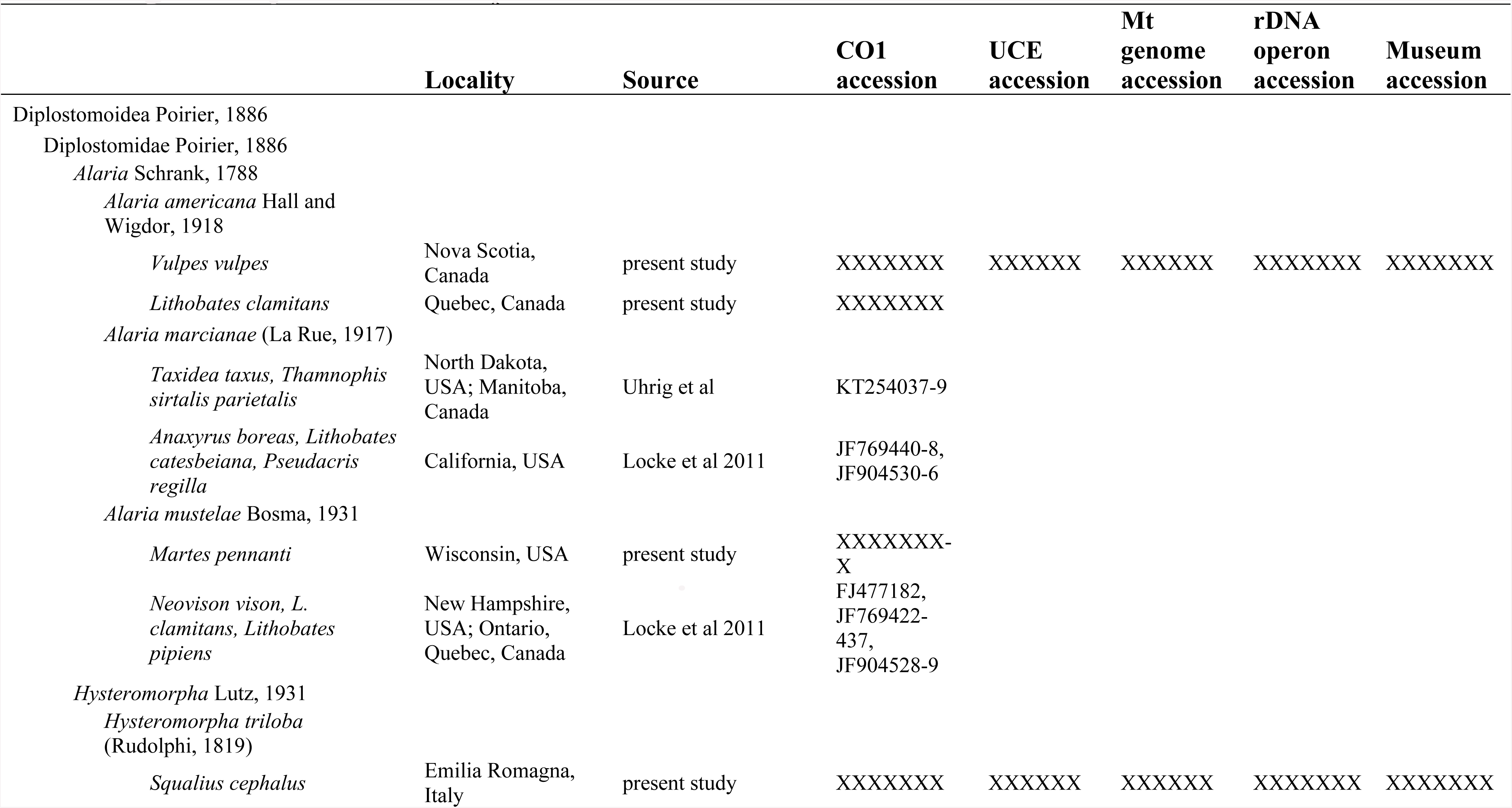
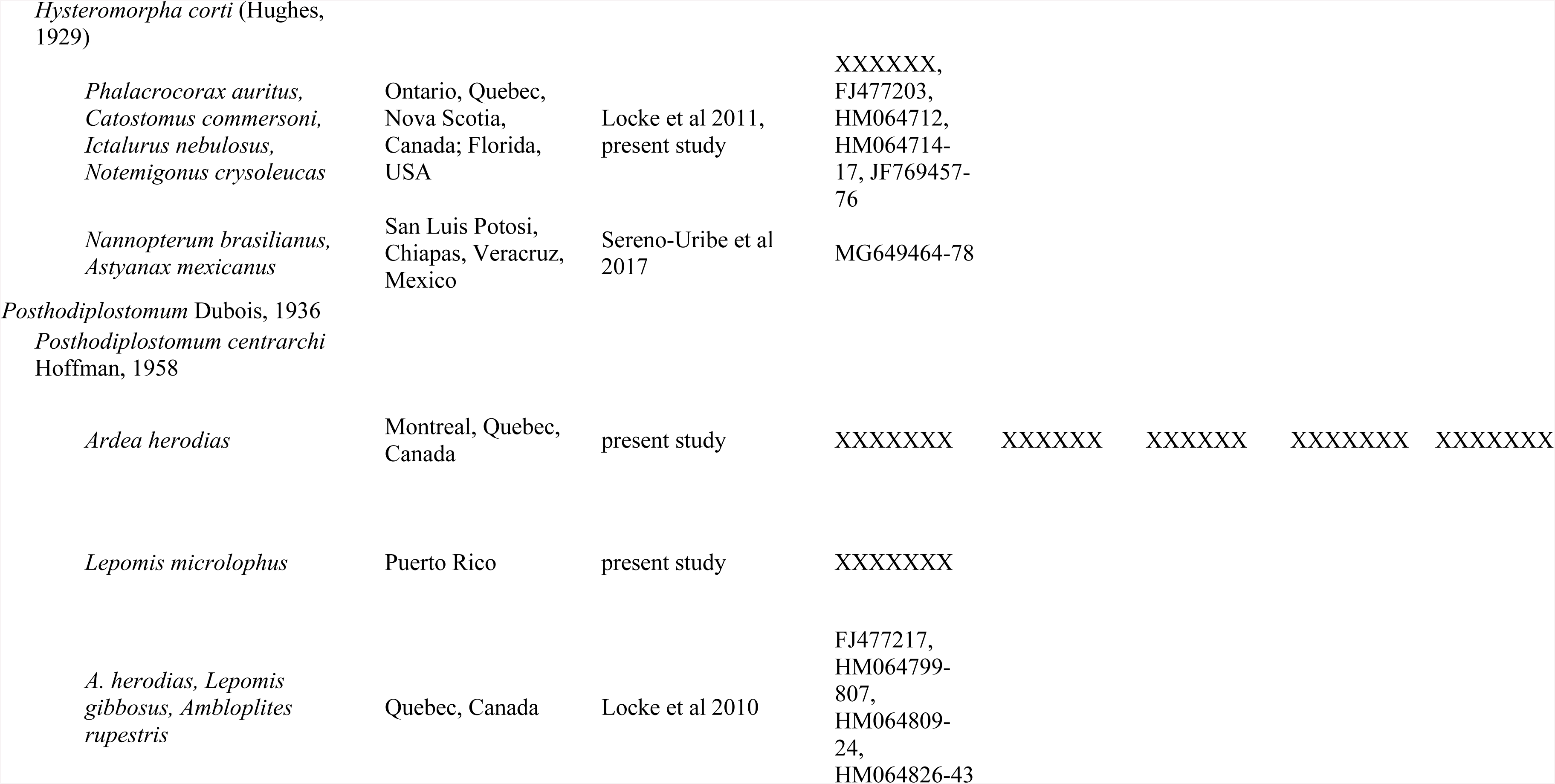
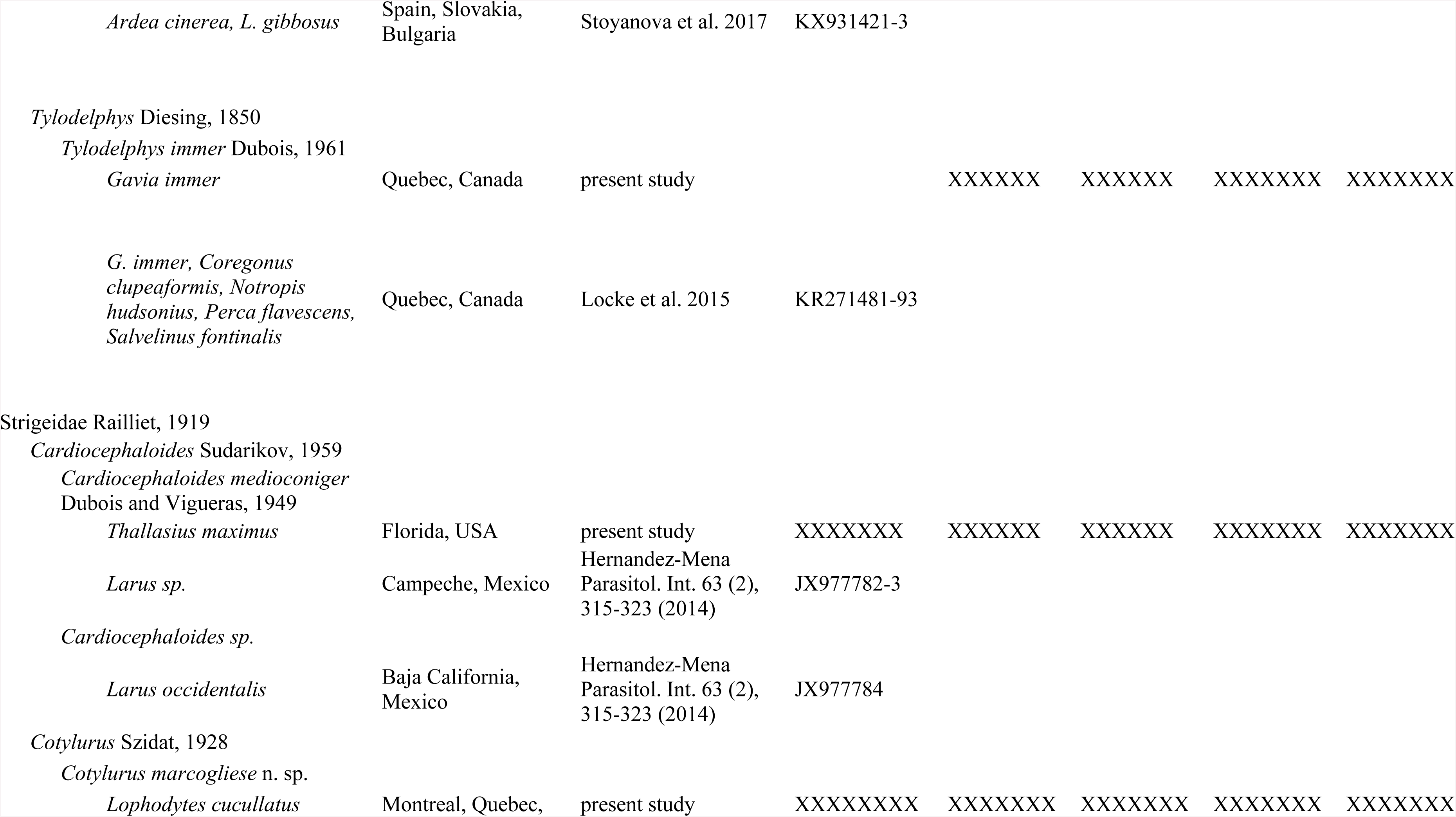
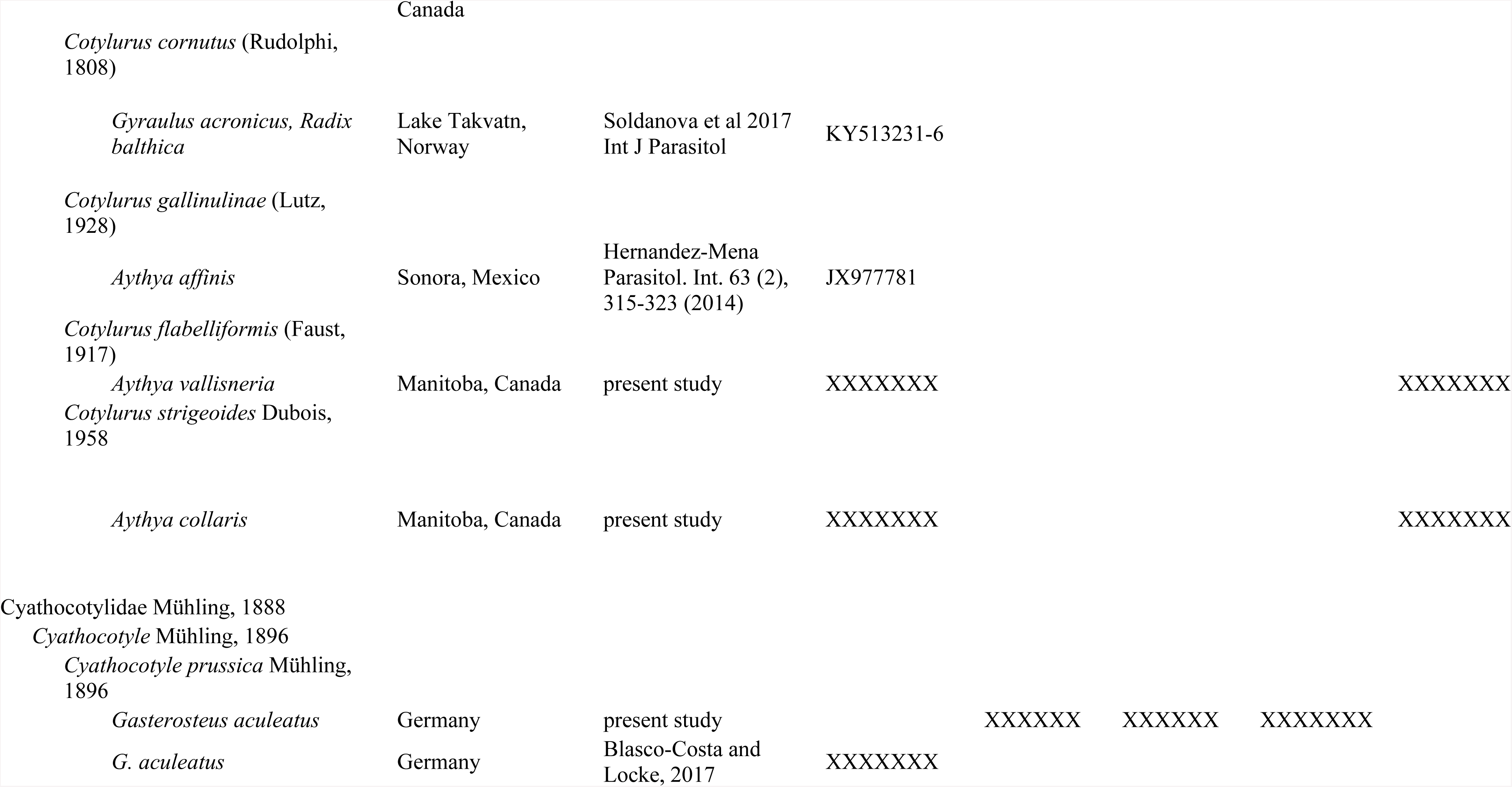
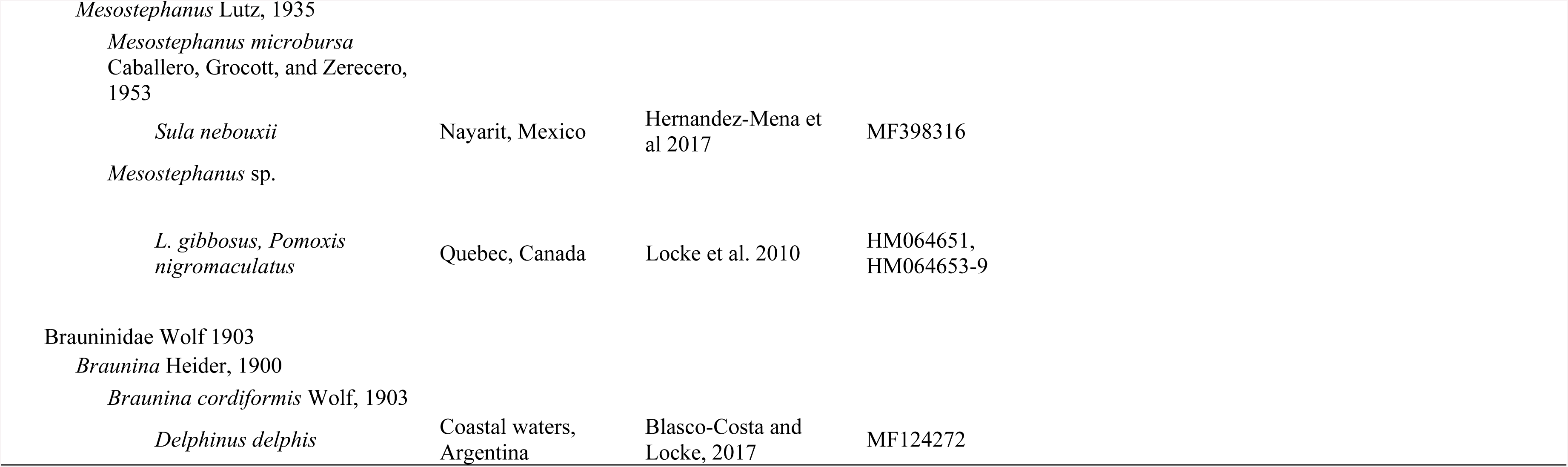
Origins of samples and data in the present and other studies

Additional conspecific or closely related parasites that contributed to identification of the seven worms, or constituted relevant host or geographic records, were processed morphologically and with PCR and Sanger sequencing (see below). Morphological vouchers were cleared in Amman’s lactophenol, rehydrated, stained in dilute acetocarmine, dehydrated in ethanol, cleared in clove oil, and mounted on a slide in Canada balsam. Vouchers comprised hologenophores (DNA sequenced from the worm studied morphologically), paragenophores (similar worms from the same individual host paired for either morphological or molecular work) and syngenophores (voucher worms from different host individuals collected at the same time, or on other occasions, or in the same region, as the worms from which DNA was extracted) (Pleijel et al., 2008). Drawings and measurements were made with the aid of a camera lucida and ocular micrometer.

### 2.2 Molecular and phylogenetic analysis

Seven samples were processed on a single lane of an Illumina HiSeq 4000 and 150-bp paired-end libraries were built using Illumina TrueSeq adapters at the UC Davis Genome Center. Partial CO1 barcodes were obtained using the primers and PCR protocols of Moszczynska et al. (2009) and Sanger sequencing, from the seven worms or other representatives, and were used to seed iterative assemblies of Illumina reads of whole mitochondrial genomes, using Geneious V9. Mitochondrial genomes were annotated in MITOS (Bernt et al., 2013) using NCBI’s Echinoderm and Flatworm Mitochondrial translation table (number 9). Additional annotations, including minor modifications to MITOS output, were made using alignments with mitochondrial genomes from two species of *Diplostomum* (Brabec et al., 2015) (KR269763-4), *Clinostomum complanatum* (KM923964), *Fasciola hepatica* (NC_002546.1), *Trichobilharia regenti* (NC_009680.1), and *Schistosoma japonicum* (NC_002544.1). A similar approach was taken to assemble complete rDNA operons (18S, ITS1, 5.8S, ITS2, and 28S). Illumina reads were mapped to rDNA sequences from, in some cases, the same samples or other representatives obtained using primers in Galazzo et al. (2002); Olson et al. (2003), and in other cases, using previously published sequences from the same species or close relatives (GB ACCESSIONS XXXXXX).

The seven mitochondrial genomes were analyzed with previously published data from Digenea and Eucestoda. Protein-coding regions were extracted and placed in the same order (i.e., atp6+nad2 and nad3 were placed before and after nad1, respectively in *Schistosoma haematobium, S. spindale*, and *S. mansoni*, in which the order of these genes differs). Protein-coding regions were concatenated, and both nucleotide and translated into amino acid sequences were aligned using MAFFT (L-INS-I). Alignments were stripped of columns with gaps and analyzed with RAXML (Silvestro and Michalak, 2012; Stamatakis, 2014) using a substitution model selected with the Bayesian Information Criterion (Tamura et al., 2013).

For UCE work, quality trimming was conducted with bbduk from the BBMap package (Bushnell, 2014). *De novo* genomes were assembled for each sample with IDBA-Hybrid (Peng et al., 2012) using *Schistosoma mansoni* (GCF_000237925.1) to help scaffold similar regions. The quality of *de novo* assemblies was assessed with BUSCO (Simão et al., 2015). Conserved genomic regions across were identified using PHYLUCE v1.6 (Faircloth, 2016). Trimmed reads were aligned to the *S. mansoni* genome using stampy-1.0 (Lunter and Goodson, 2011) with a substitution rate set to 0.05 to capture overlapping regions with trimmed data. Using overlapping regions from the *S. mansoni* genome, an initial probe set was mapped back to the *de novo* genomes via PHYLUCE scripts. Genomes were similarly analyzed from *Clonorchis sinensis* (PRJDA72781), *Echinostoma caproni* (PRJEB1207), *Diphyllobothrium latum* (PRJEB1206), and *Fasciola hepatica* (PRJEB6687).

Complete probe sets were generated for ten out of twelve species (phyluce_probe_query_multi_fasta_table in PHYLUCE), with 801 loci and 16,671 3 ×; tiling probes post-duplicate-filtering. Loci were aligned using MAFFT and with 90% matrix occupancy, 517 loci were retained. The MAFFT-Gblocks option was called in PHYLUCE to remove poorly aligned regions. Phylogenetic reconstructions were computed using RAxML (Stamatakis, 2014) from a concatenated matrix with GTRGAMMA substitution rate, with 1,000 bootstrap replicates in each of 20 parallel runs, and *Diphyllobothrium latum* set as outgroup. The resulting maximum-likelihood (ML) tree with bootstrap bi-partitions was annotated with morphological and life-history characters from the Diplostomoidea (supplementary Fig. 1). We also generated one tree without setting *D. latum* as the outgroup, and another after running PHYLUCE and RAxML solely with the seven diplostomoids and *S. mansoni* set as outgroup (the latter yielding 796 conserved genetic elements, and 63,959 variable sites based on 85% matrix occupancy). In both the latter cases (not shown), the topology obtained was well supported and indistinguishable from the UCE tree reported below.

## 3. Results

### 3.1 General molecular results

The genomic analysis of seven samples yielded 7.8×10^8^ 150-bp reads, mean 1.1×10^8^, range 2.1×10^7^–1.4×10^8^ reads per sample; mean N50=2833 bp, range 542-11122 bp). Below, and in Table 2, these and other results are broken down by species.

**Table 2:** Selected assembly statistics for 150-bp paired-end reads from IDBA_hybrid.

|  | **Reads** | **Aligned**<br>**reads** | **Expected**<br>**coverage** | **Contigs** | **N50** | **max** | **mean** | **Total length** | **N80** |
| --- | --- | --- | --- | --- | --- | --- | --- | --- | --- |
| Cyathocotylidae |  |  |  |  |  |  |  |  |  |
| *Cyathocotyle*<br>*prussica* | 21208232 | 7029966 | 0.055976 | 336463 | 542 | 13687 | 522 | 175748574 | 396 |
| Strigeidae |  |  |  |  |  |  |  |  |  |
| *Cardiocephaloides*<br>*medioconiger* | 91513844 | 72475903 | 0.120399 | 566427 | 11122 | 157631 | 1204 | 682202373 | 2185 |
| *Cotylurus*<br>*marcogliesei* n. sp. | 136959212 | 80825403 | 0.133569 | 2065313 | 1662 | 217472 | 427 | 883621846 | 204 |
| Diplostomidae |  |  |  |  |  |  |  |  |  |
| *Alaria americana* | 130527714 | 71565054 | 0.072403 | 1899210 | 1540 | 90097 | 658 | 1249851331 | 507 |
| *Hysteromorpha*<br>*triloba* | 130953814 | 78792229 | 0.076501 | 1923782 | 1675 | 89784 | 675 | 1299457195 | 496 |
| *Posthodiplostomum*<br>*centrarchi* | 143789074 | 81923449 | 0.105401 | 2157712 | 1245 | 878066 | 497 | 1074511666 | 329 |
| *Tylodelphys immer* | 123480600 | 80297700 | 0.155342 | 1736417 | 2044 | 382985 | 432 | 751015059 | 201 |

### 3.2 Phylogenetic analysis of mitochondrial genomes

The mitochondrial genomes were 13,665 – 15,107 bp in length (supplementary Table 1). The 12 coding regions occurred in the same order as in digeneans other than *S. haematobium, S. spindale*, and *S. mansoni*. In *C. prussica*, two pairs of tRNA genes were reversed in order compared with other diplostomoids. One of these reversals, the occurrence of trnN prior to trnP, is also a feature of *Clinostomum complanatum* (KM923964). The order of three tRNA genes occurring between *nad6* and *nad5* in *C. medioconiger* differed from that seen in other diplostomoids, and these genes were positioned adjacent to a 774-bp non-coding span which, among diplostomoids and *Clinostomum*, was unique to *C. medioconiger*.

Phylogenetic analysis of nucleotide or translated amino acid sequences from the 12 concatenated coding regions of the mitochondrial genomes did not support the order Diplostomida (Fig. 2). The diplostomoids and *Clinostomum* emerged as early diverging descendants from common ancestors of the Plagiorchiida, rather than grouping with *Schistosoma* and *Trichobilharzia*.

### 3.3 Phylogenetic analysis of ultra-conserved elements (UCEs)

The Diplostomoidea were monophyletic in the topology obtained from ML analysis of 517 conserved genetic elements, comprising 84,902 distinct patterns across 234,783 characters per taxon (Fig. 3). Consistent with the concept of the Diplostomida and Plagiorchiida, the Diplostomoidea and *Schistosoma* formed a clade separate from a clade comprising plagiorchiidans *Clonorchis, Echinostoma*, and *Fasciola*.

Relationships among diplostomoids were similar to those recovered by analysis of mt genomes, differing slightly within a clade of Diplostominae *Alaria* + *Hysteromorpha* + *Tylodelphys* + *Diplostomum* (Figs. 2, 3). Within the diplostomoids, *Cyathocotyle prussica* was basal to a clade in which the Diplostomidae were paraphyletic. The diplostomid *Posthodiplostomum centrarchi* was basal to a clade of both strigeids and other diplostomids. The two strigeids were sister taxa.

The non-molecular characters of metacercariae showed higher correspondence with the UCE topology than did the characters of adults (Fig 3, supplementary Fig. 1). Within the Diplostomoidea all five morphological or life-history characters of metacercariae mapped onto genomic clades, while correspondence was poorer for adult characters. Metacercarial apomorphic characters included the structure of the reserve bladder, presence or absence of encystment, free or enclosed limebodies, and the presence or absence of pseudosuckers, which distinguished clades comprising relatively small subsets of species. For example, the structure of the reserve bladder differs among the following four groups, which correspond to clades in the molecular phylogeny: *Cyathocotyle, Posthodiplostomum, Cardiocephaloides* + *Cotylurus*, and *Alaria* + *Hysteromorpha* + *Tylodelphys*. Commonly recognized metacercarial morphotypes (prohemistomulum, neascus, tetracotyle, diplostomulum) mapped onto all clades formed.

### 3.4 Morphological and molecular support of identification or description of species

Specimens were identified morphologically, and dimensions are reported in μm and given as range (means, ±standard deviation, n measured). Sequences of partial sequences of cytochrome *c* oxidase 1 (DNA barcodes) from mitochondrial genomes were also compared to previously published data and with sequences from additional collections. In a neighbour-joining phenogram (Fig. 4), CO1 sequences fell into distinct clusters in which the minimum p-distance between nearest neighbour species was in all cases less than the maximum distance within species; these clusters formed well supported clades in separate ML analysis. Complete rDNA operons (Supplementary Table 1) were subjected to BLAST searches, which supported identifications in all cases, as described further below.

**Fig 4.**
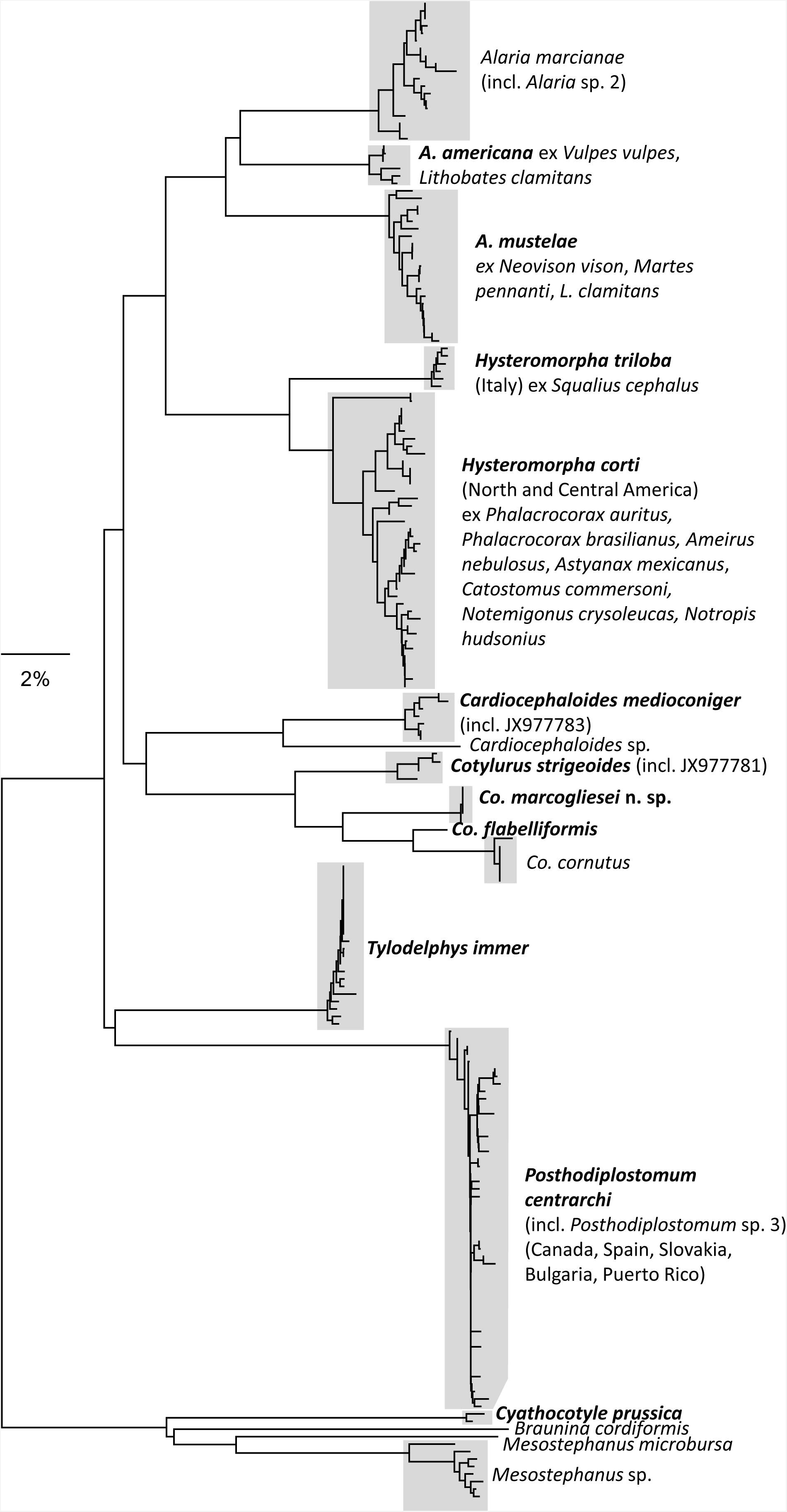
Neighbour-joining tree of uncorrected p-distance among 195 partial sequences of cytochrome *c* oxidase I (CO1). Shaded clades had >99% support in 500 bootstrap replicates in ML analysis (not shown) and are annotated with identifications, host and geographic records mentioned in the results. Sequences from the present study were obtained from species labelled in bold. This includes Sanger-sequenced amplicons of the barcode region of COI (n=35, XXXXXX-X) and mitochondrial genomes of specimens used in Figures 2 and 3. Sequences (n=160) from other studies are FJ477182, FJ477203, FJ477217, HM064651-9, HM064712, HM064714-7, HM064799-843, JF769422-76, JF904528-36, JX977781-4, KR271481-93, KT254037-9, KX931421-3, KY513231-6, MF124272-3, MF398316, MG649464-78.

**Fig 5.**
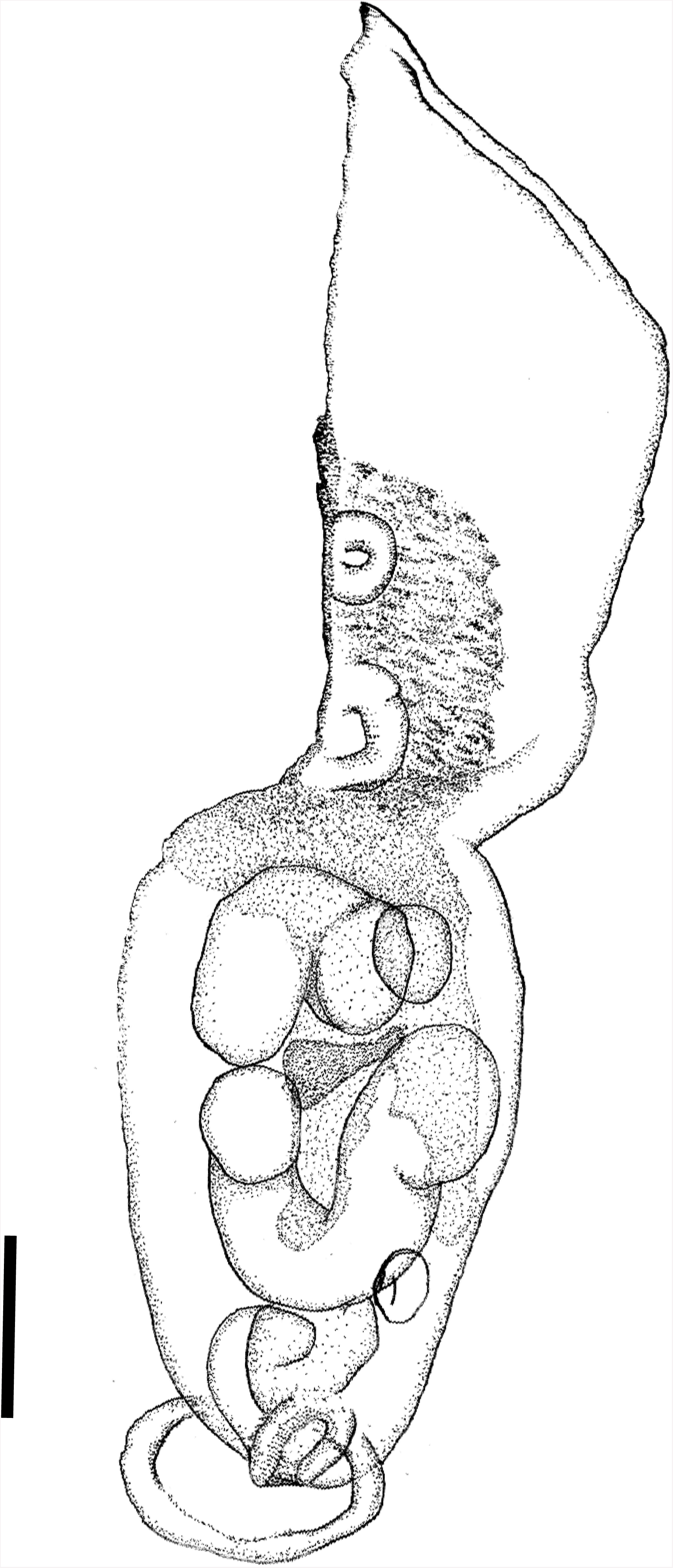
Adult of *Posthodiplostomum centrarchi* Hoffman, 1958, from *Ardea herodias*, Ile aux Herons, St. Lawrence River, Quebec, Canada. Scale = 200 μm. Hologenophore for partial sequence of cytochrome *c* oxidase I, Genbank accession XZXXXXXX.

#### 3.4.1 *Cyathocotyle prussica* Mühling, 1896

*Cyathocotyle prussica* was identified based on morphology of the unstained specimen prior to DNA extraction, and its provenance (host, geographic origin). Metacercariae of *C. prussica* have been reported from *Gasterosteus aculeatus* and other small-bodied fishes throughout Europe (Kalbe et al., 2016; Kvach et al., 2016). The DNA used for genomic work was obtained from a metacercaria from *Gasterosteus aculeatus* collected at the same time as the specimen sequenced by Blasco-Costa and Locke (2017) and no morphological voucher remains. The CO1 region of the mitochondrial genome assembly was 99.7 % (587 / 589 identities) similar to MF124273, and the rDNA operon 99.1 % (762/769 identities) similar to KY951726, both from Blasco-Costa and Locke (2017). The rDNA operon differed by 2.5-20.8% from 13 published rDNA sequences from cyathocotylids on GenBank.

#### 3.4.2 *Posthodiplostomum centrarchi* (Hoffman, 1958) Stoyanov, Georgieva, Pankov, Kudlai, Kostadinova, and Georgiev, 2017 (Fig. 5)

[Measurements from 9 specimens (6 paragenophores, 3 hologenophores) ex *Ardea herodias*, Montreal, Quebec, Canada]

Total length 1222–1775; forebody and hindbody separated by marked constriction. Forebody spatulate 680–1200 long, 452–875 wide, oval to lanceolate, with anterior lateral edges often recurved ventrally. Hindbody oval, widest in anterior half, tapering posteriorly, 520–875 long, 248–750 wide. Hindbody length / forebody length 0.58–0.86. Oral sucker terminal, 38–72 ×; 38–72. Ventral sucker 55–90 ×; 55–80, in posterior half of forebody, centered 54–71 % along forebody sagittal axis. Tribocytic organ 140–200 ×; 144–256. Pharynx 40–53 long, 40–53 wide. Oesophagus 60–72 long. Ovary lateral to anterior testis at anterior margin of hindbody, 80–105 ×; 72–88. Testes occupying first 55–80% of hindbody. Anterior testis irregularly, smoothly lobed 96–200 long, 144–272 wide. Posterior testis, sinuous, v-shaped, pointed posteriorly, 176–350 long, 216–350 wide. Vitelline reservoir intertesticular, median. Vitellaria densest at level of posterior of tribocytic organ and in base of forebody, extending anteriorly beyond ventral sucker by slightly more than one ventral-sucker length, in hindbody extending more than halfway to posterior extremity and bifurcating into two posteriorly oriented bands at and beyond vitelline reservoir, sometimes exceeding, sometimes exceeded by, posterior extent of posterior testis. Copulatory bursa eversible, terminal or slightly dorsal, 120–216 long, 135–300 wide, housing muscular genital cone. Eggs few, 0–4, length 70–98 ×; 42–64 (See supplementary Table 2 for means, standard deviations, n structures measured).

##### 3.4.2.1 Remarks

MacCallum (1921) summarily described *Posthodiplostomum* (*Diplostomum*) *minimum* from *Ardea herodias* in New York, USA. Hoffman (1958) created two sub-species, *P. minimum centrarchi* and *P. minimum minimum*, for lineages infecting either centrarchids or cyprinids, respectively. Stoyanov et al. (2017) elevated the centrarchid subspecies to *P. centrarchi.*

Our observations largely agree with Dubois’ (1970b) description of the adult (although he did not distinguish *P. centrarchi* and *P. minimum*), except in the following respects (Supplementary Table 2): Two of nine specimens were up to 75 greater in total length; in 4/9 specimens, we observed the hindbody to be widest at the anterior testis (e.g., Fig 5), rather than at the level of the posterior testis as in Dubois (1970b). The oral sucker was the same length as the pharynx in 1/6 specimens, rather than larger as in Dubois (1970b); the tribocytic organ was wider in 3/6 specimens (200-256 *versus* maximum of 190 in Dubois, 1970b); and in 7/9 specimens the copulatory bursa was both longer and wider than maximum values (160 ×; 160) reported by Dubois (1970b). The dimensions of the specimens we encountered also exceed values reported by Palmieri (1977), who reported means and standard deviations from adults from diverse hosts experimentally infected with metacercariae from centrarchids. However, as seen above, morphometric deviations relative to these studies were small, and Palmieri (1977) showed that adult morphology varies a great deal within *P. centrarchi*. Also, deviations were obtained from hologenophores genetically matching *P. centrarchi* common in centrarchid hosts (e.g., see records in (Boone et al., 2018; Locke et al., 2010)), and therefore there is no doubt they represent *P. centrarchi*.

The worm used in genomic analysis had partial CO1 sequences and ITS1-5.8S-ITS2 sequences 99-100% similar to those of *P. centrarchi* (=*Posthodiplostomum* sp. 3 of Locke et al., 2010; Stoyanov et al., 2017), including the hologenophore in Fig. 5, and CO1 from *P. centrarchi* from liver of *Lepomis microlophus* in Puerto Rico.

#### 3.4.3 *Cardiocephaloides medioconiger* Dubois and Vigueras, 1949 (Fig. 6)

[Measurements from 3 hologenophores ex *Thallasius maximus*, Florida Keys, USA]

**Fig 6.**
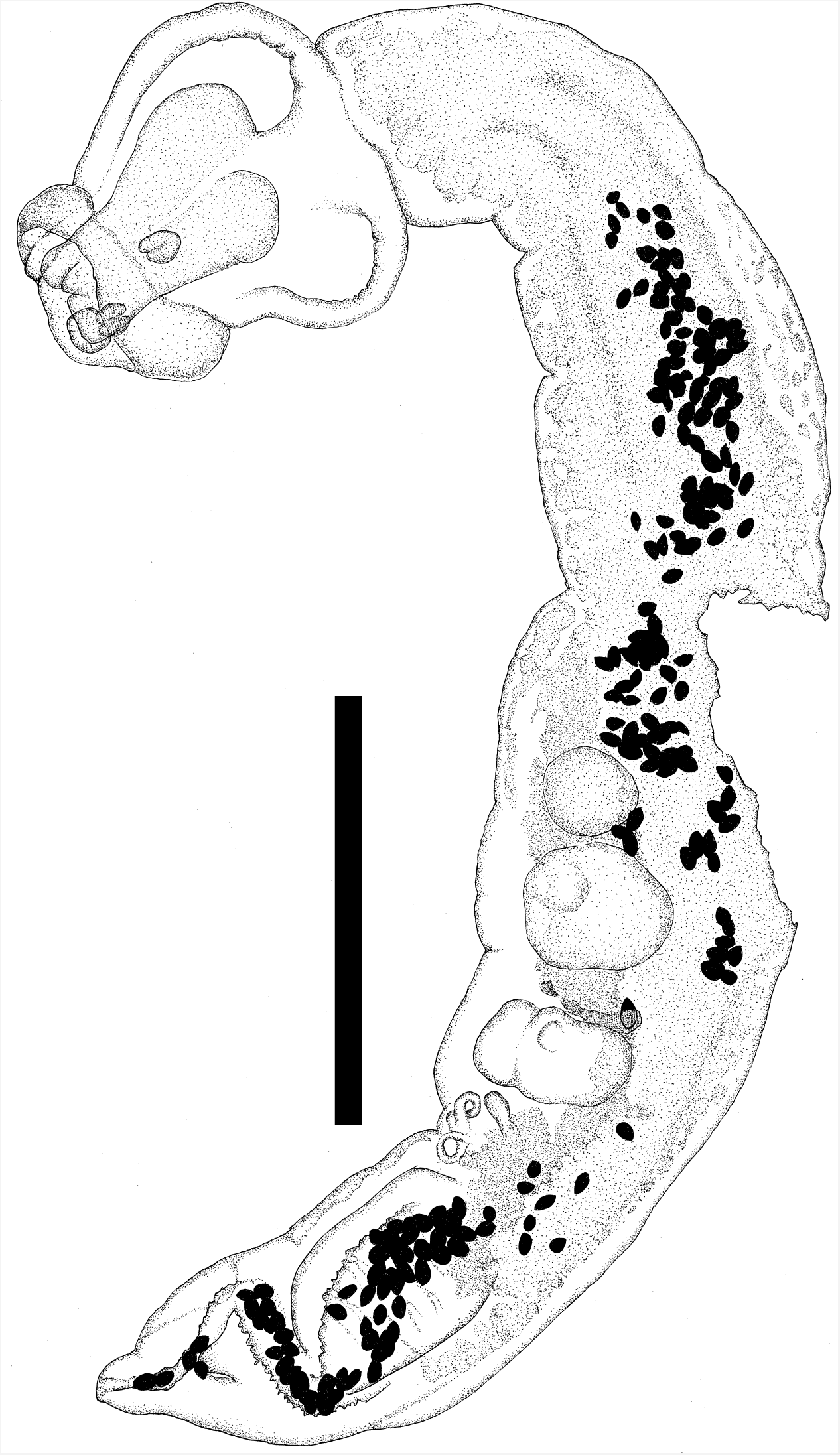
Adult of *Cardiocephaloides medioconiger* Hall & Wigdor, 1918, from *Thalassius maximus*, Florida, United States. Scale = 2000 μm. Hologenophore for partial sequence of cytochrome *c* oxidase I, Genbank accession XZXXXXXX.

Total length 7273–8324; forebody and hindbody separated by moderate constriction. Forebody tulip-shaped 1333–1616 long, 1414–1455 wide. Hindbody 5657–6869 long, 1232–1293 wide, width gradually increasing posteriorly, widest at level of testes, tapering to point at posterior extremity. Hindbody 3.5–4.9 as long as forebody. Oral sucker 103–160 ×; 175–193. Ventral sucker 103–168 ×; 112–128. Tribocytic organ bi-lobed, with one lobe well developed and darkly staining. Pharynx 129–152 long, 119–152 wide. Ovary pretesticular, 363–363 long, 300–363 wide. Vitellaria dense in anterior hindbody, absent from ventral surface in region of ovary and testes, extending dorsally along anterior of copulatory bursa. Testicular zone 828–1010 long. Anterior testis 303–475 long, 606 wide. Posterior testis 363–484 long, 485–707 wide. Vitelline reservoir intertesticular. Eggs numerous 94–117 long, 62–70 wide with shells 2.0–2.2 thick. Copulatory bursa 1010–1919 long. Hindbody length 3.6–5.6 times that of copulatory bursa (means, standard deviations, n structures measured in Supplementary Table 3).

##### 3.4.3.1 Remarks

*Cardiocephaloides medioconiger* was described from *Larus argentatus* in Cuba and has been reported from *T. maximus* in the same region (Dubois, 1970b). The morphology of the three voucher specimens was consistent with Dubois (1970b) (Supplementary Table 3), except that the following were larger: forebody (maxima of length and width of 1500 and 1360, respectively, in Dubois, 1970b), oral sucker (maximum width 136 in Dubois, 1970b) and ovary (maximum 278 ×; 300 in Dubois, 1970b). The four CO1 barcode sequences obtained were 99.1-99.8% similar to *C. medioconiger* (JX977783) from *Larus* sp. collected in Campeche, Mexico (Hernández-Mena et al., 2014). Within species of *Cardiocephaloides*, mean variation in CO1 is 0.7% (range 0-1.5%) and between species, 8.8% (range 7.4 −9.7%). The rDNA operon from the specimen we collected differed by 1.2% (1669/1672 identities) from 18S (MF398359, isolate DNA181) and 2.3% from the ITS (1041/1065 identities with JX977844, isolate DNA181) of *Cardiocephaloides* sp. from *Larus occidentalis* in Baja California, Mexico, and by 0.5% (1839/1848 identities) from 18S from *C. longicollis* (AY222089) from *Larus ridibundus*, Ukraine.

##### 3.4.4.1 *Cotylurus marcogliesei* n. sp. (Fig. 7)

##### 3.4.4.2 Description

[Measurements from 11 specimens (3 hologenophores, 7 paragenophores, and holotype), ex *Lophodytes cucullatus*]

**Fig 7.**
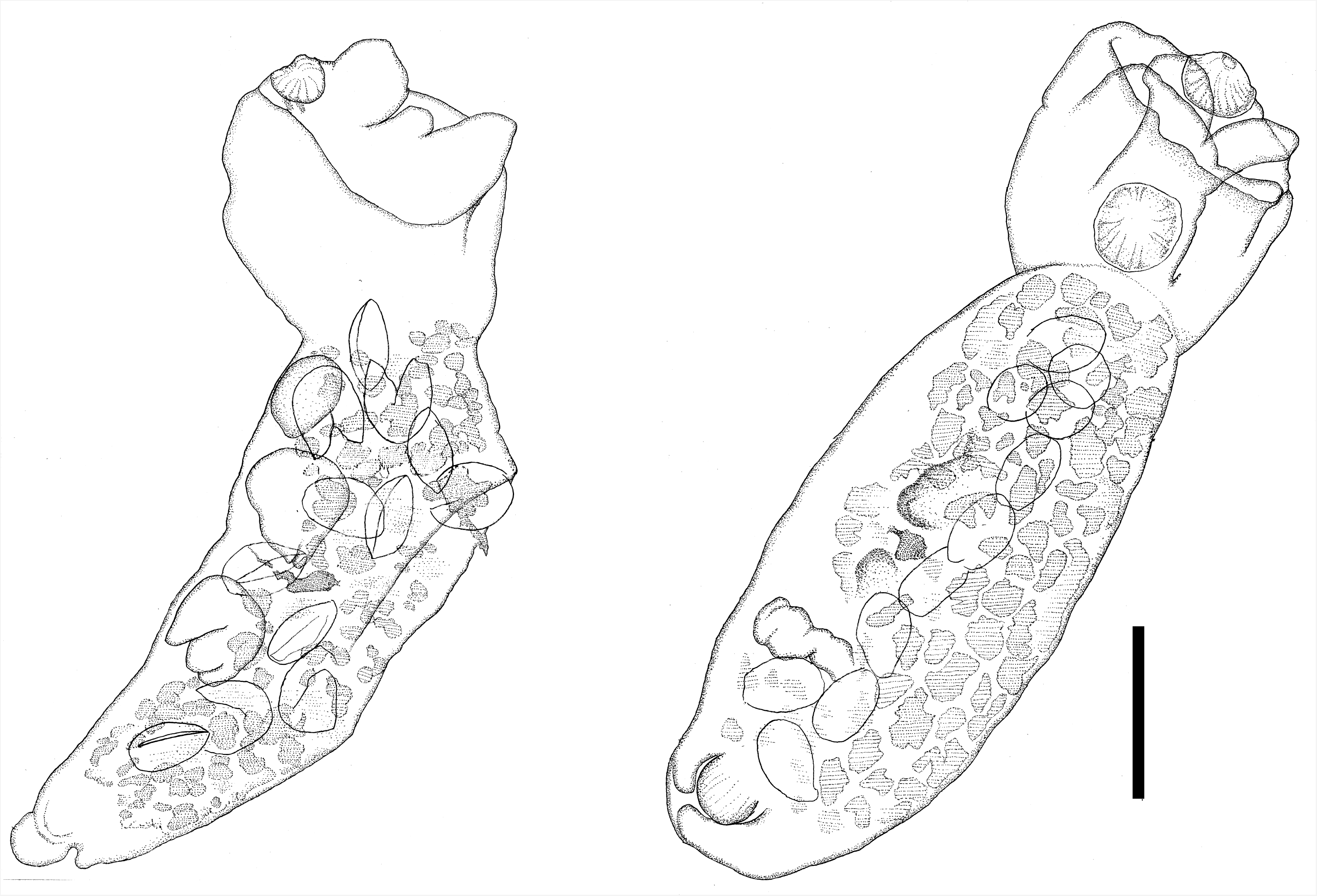
Adults of *Cotylurus marcogliesei* n. sp. from *Lophodytes cucullatus* in Montreal, Quebec, Canada. Scale = 200 μm. Hologenophores for Genbank accession XZXXXXX.

Adult total length 816–1152 (973±107, 10); forebody and hindbody separated by marked constriction. Body mildly arched dorsally. Forebody cup-shaped 216–408 (307±51, 11) long, 312–640 (436±95, 10) wide, with broad, oblique opening. Ventral forebody wall markedly shorter than dorsal. Hindbody stout along entire length, 600–880 (703±96, 10) long, 256–520 (318±81, 9) wide. Forebody to hindbody length ratio 1:1.9–2.8 (2.3±0.3, 10). Oral sucker terminal, 48–80 (68±13, 9) ×; 64–112 (85±15, 8). Ventral sucker 64–128 (99±24, 6) ×; 80–168 (108±28, 7). Tribocytic organ bilobed, with one or both lobes extending anteriorly beyond margin of forebody; proteolytic gland not observed. Pharynx small, difficult to observe, 28–34 (30±3, 3) long, 33–56 (43±12, 3) wide. Testes tandem, small, lobed and smooth; anterior testis 112–168 (132±27, 4) long, 112–120 (116±6, 2) wide, posterior margin at 32–45 (38±6, 4) % of hindbody. Posterior testis 136–160 (144±11, 4) long, 104–116 (110±8, 2) wide, with posterior margin at 59–67 (64±3, 5) % of hindbody. Ovary, near anterior extremity of hindbody, oval, 40– 96 (68±40, 2) long, 56–56 (56±0, 2) wide. Vitellaria follicular, confined to hindbody, densely distributed in ventro-lateral field extending posteriorly to level of copulatory bursa and genital bulb, without obscuring the latter. Vitelline reservoir intertesticular; median. Uterus with 4–13 (9±3, 9) eggs, 76–108 (95±7.6, 37) long ×; 40–72 (56±6.9, 37) wide. Copulatory bursa large, genital bulb round, oval or reniform, 72–144 (111±27, 7) ×; 74–160 (117±27, 7).

##### 3.4.4.3 Diagnosis

Adults of *Cotylurus marcogliesei* n. sp. possess a typically strigeid morphology, vitellaria confined to the hindbody, genital bulb in the copulatory bursa and smooth, bi- or trilobed testes with lobes pointing posteriorly, all of which are characteristic of *Cotylurus*. The wide opening of the forebody is more representative of *Ichthyocotylurus* Odening, 1969 (Niewiadomska, 2002b), but the presence of a genital bulb and testes with posterior facing lobes, in addition to CO1 sequence similarity (Fig. 4) indicate the species belongs to *Cotylurus*. The most morphologically similar species is *C. brevis* Dubois and Rausch, 1950, from which molecular data are unavailable. The hindbody of *C. marcogliesei* n. sp. is 2.1–2.8 times as long as the forebody, while in *C. brevis* it is 1.3–1.9 times as long (Dubois, 1970b) (Supplementary Table 4). Mature adults of *C. marcogliesei* n. sp. are 816–1152 (mean 990) in total length, and half of the specimens were shorter than the minimum length of *C. brevis* (1000–1800) (Dubois, 1970b). The pharynx is shorter in *C. marcogliesei* n. sp. (28-34) than in *C. brevis* (50-59), although this organ is difficult to visualize in both species (Dubois, 1970b) and may not be a reliable character for identification or discrimination. To our knowledge, no members of the genus *Cotylurus* have been recorded from *Lophodytes* (Anatidae, Merginae), which further distinguishes it from *C. brevis*, originally described in Europe and found mainly in *Aythya* and *Anas* spp. (Anatidae, Anatinae).

> Type host: *Lophodytes cucullatus* (definitive host)
>
> Site of infection: Small intestine (definitive host)
>
> Type locality: Montreal, Quebec, Canada (50.183 N, –98.383 W) (definitive host)
>
> Type material: Holotype (adult worm) Voucher accessions forthcoming;
>
> Representative DNA sequences: CO1: XXXXXXX.
>
> Etymology: The species is named after David J. Marcogliese, for his contributions to parasitology.

Partial CO1 sequence was obtained from one of two specimens of *C. flabelliformis* from *Aythya vallisneria* collected in Manitoba. The paragenophore was 782 long, with forebody 351 ×; 367, strongly arched dorsally, hindbody 638 ×; 399, oral sucker 64 ×; 96, pharynx 44 ×; 44, ventral sucker 88 ×; 112, eggs (n=30) 93-100 ×; 50-60, and its dimensions, morphology, host and geographic provenance agree with the collective accounts of Dubois (1970b), Campbell (1971) and Lapage (1961). Sequences of CO1 were obtained from three specimens of *Cotylurus strigeoides* Dubois, 1958 from *Aythya collaris*. The three hologenophores were 1818 long, with forebody 64 long, hindbody 1313-1475 ×;747-768, oral sucker 160 ×; 112, pharynx 88 ×; 76, ventral sucker 192 ×; 136, ovary 152-231×128-207, anterior testis 223-283 ×; 239-423, posterior testis 271-319 ×; 271-343, eggs (39≤n≤79) 88-98 ×; 56-64. The CO1 sequences from *C. strigeoides* were 1.1-1.5% divergent from a CO1 sequence (JX977781) of a worm from *Aythya affinis* collected in Sonora, Mexico (Hernández-Mena et al., 2014). Because of the low level of divergence between *C. strigeoides* and JX977781, we believe all these data originate from *C. strigeoides*, but JX977781 is identified as *C. gallinulinae*. The material we examined was distinguished from *C. gallinulinae* by a large pharynx (5% of total length, and over half the size of the oral sucker, versus ≤2% of total length, and less than half the size of the oral sucker in *C. gallinulinae*, Dubois, 1970b), and the position of the ovary immediately posterior to the division of the fore- and hindbody (Dubois, 1958), whereas in *C. gallinulinae* it lies further posterior (Fig. 211 in Dubois (1970b). Records in Manitoba and Sonora are plausible for *C. strigeoides*, which is known from California and Alaska, whereas *C. gallinulinae* is neotropical (Dubois, 1970b; McDonald, 1981). Both JX977781 and the specimens we identified as *C. strigeoides* were from anatid hosts, which is typical for *C. strigeoides*, whereas *C. gallinulinae* is known from members of the Raillidae (Dubois, 1970b; McDonald, 1981). Within *Cotylurus* as whole, CO1 varies 0-0.3% (mean 0.2%) within species and 3.4-11.2% (mean 8.3%) between species (considering JX977781 as *C. strigeoides*). Sequences of CO1 from *C. marcogliesei* n. sp. differed by 7.9-9.3% from other species of *Cotylurus*. The rDNA operon of *C. marcogliesei* n. sp. differs from other species of *Cotylurus* by 0.7-3% (JX977841, KY513180-2, MF398340).

#### 3.4.5 *Alaria americana* Hall and Wigdor, 1918 (Fig. 8)

[Measurements from 8 specimens (2 hologenophores, 6 paragenophores), ex *Vulpes vulpes*, Nova Scotia, Canada]

**Fig 8.**
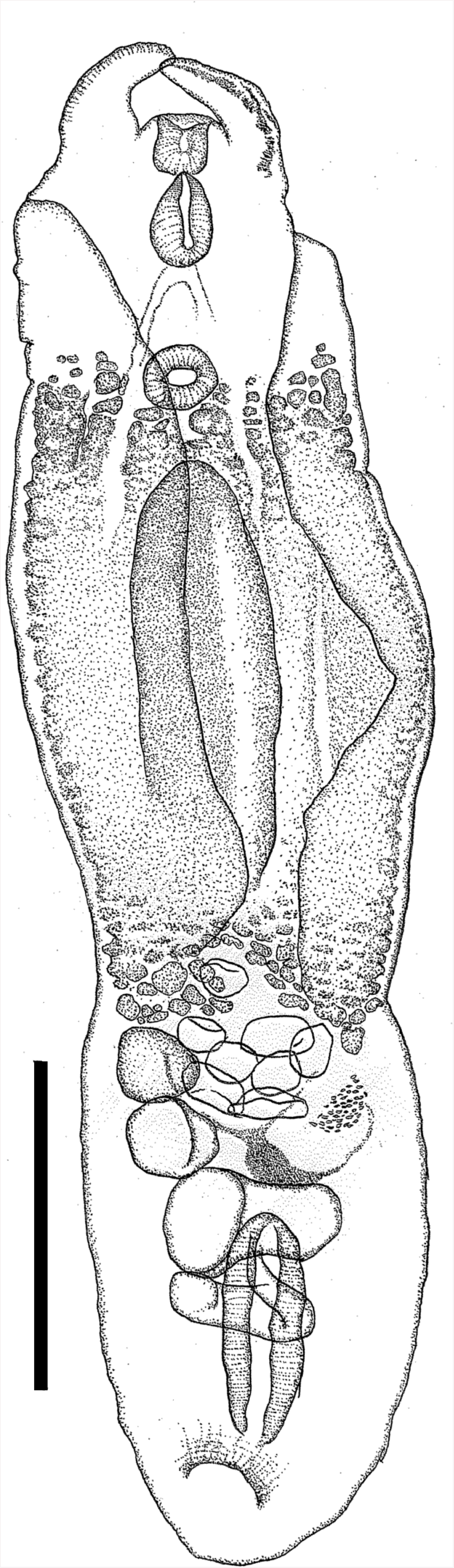
Adult of *Alaria americana* Hall & Wigdor, 1918, from *Vulpes vulpes*, Nova Scotia, Canada. Scale = 500 μm. Paragenophore for partial sequence of cytochrome *c* oxidase I, Genbank accession XZXXXXXX.

**Fig 9.**
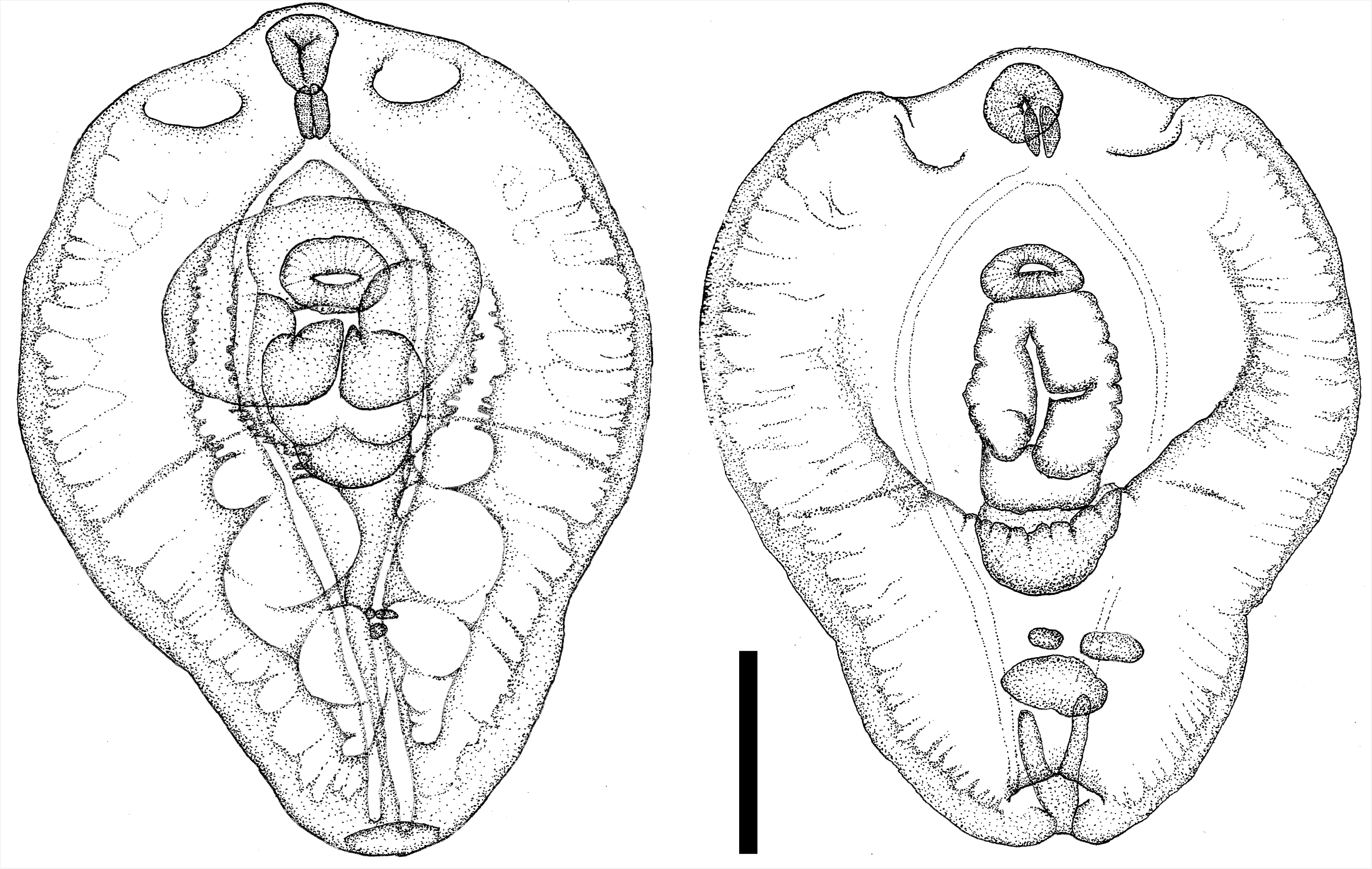
Metacercariae of *Hysteromorpha triloba* (Rudolphi, 1819) from *Squalius cephalus*, Italy. Scale = 200 μm. Paragenophores for partial sequence of cytochrome *c* oxidase I, Genbank accession XZXXXXXX.

Total length 2121–2868; forebody and hindbody separated by shallow constriction. Forebody with foliaceous lateral margins folded over ventral surface, 1375–1858 long, 465–869 wide. Hindbody oval, 606–1010 long, 252–559 wide. Forebody 1.6–2.3 times longer than hindbody. Lappets 136–240 long, with stippled glandular tissue along outer margin and protruding from anterior extremity lateral to oral sucker, 64–120 ×; 50–119. Ventral sucker 88–104 ×; 88–120. Tribocytic organ originating posterior to ventral sucker, 667–788 long, 160–250 wide. Pharynx muscular, pyriform, 120–150 long, 45–96 wide, wider posteriorly, larger than oral sucker. Pharynx length 1.2–2.0 times oral sucker length. Testes in anterior two thirds of hindbody. Anterior testis smooth, unevenly lobed, lateral, 110–319 ×; 120–327, situated opposite Mehlis’ gland. Posterior testis sub-symmetrical, extending laterally across hindbody width, with two lateral, round lobes, 80–223 ×; 152–270. Ovary lateral near anterior extremity of hindbody, round to oval, 67–160 ×; 65–280 wide. Vitellaria mainly in forebody, dense in tribocytic organ, extending anterior to or just past ventral sucker, extending posterior to or slightly beyond forebody-hindbody constriction. Ejaculatory pouch fusiform, 250–444 ×; 101–135, with muscular walls 31–55 thick, extending from posterior testis, or just posterior, to dorsally opening genital atrium. Seminal vesicle with sinuous with transverse sections lying dorsal to ejaculatory pouch, posterior to posterior testis. Vitelline reservoir intertesticular; median. Eggs 0–11, 102–136 long ×; 36–80 wide. (means, standard deviations, n structures measured in Supplementary Table 5)

##### 3.4.5.1 Remarks

Dubois (1970b) considered *A. americana* a junior synonym of *A. marcianae*. The morphological, molecular and life-history data herein support Johnson (1970) and Pearson and Johnson (1988), who maintained *A. americana* as valid (Supplementary Table 5). Adults of *A. americana* are larger (>2000) than those of *A. marcianae* (<2000) and have a thicker-walled ejaculatory pouch (>20 compared with <20 in *A. marcianae*). The CO1 from adults from *Vulpes vulpes* in Nova Scotia matched (i.e., 98.4-99.8% similarity) sequences from two mesocercariae from a *Lithobates clamitans* in Quebec. Both *V. vulpes* and *L. clamitans* are known hosts of *A. americana* in North America (Dubois, 1970b). Adults of this species occur in canid definitive hosts, while *A. marcianae* mainly matures in felid and mustelid hosts (Pearson and Johnson, 1988). The adult *A. marcianae* sequenced in Uhrig et al. (2015) was collected from *Taxidea taxus* (Mustelidae).

In *A. americana* and other *Alaria* spp., intraspecific CO1 distances range from 0 to 2.9% and interspecific distances from 8.4 to 13.1% (total 53 CO1 sequences from *A. americana, A. mustelae*, and *A. marcianae; A. marcianae* includes *Alaria* sp. 2 of Locke et al. (2011) as noted by Uhrig et al. (2015), who recorded the sequence match but mistakenly referred to *Alaria* sp. 1 of Locke et al. (2011)). We also include here new CO1 records from *A. mustelae* from *Martes pennanti* from Wisconsin.

The rDNA operon of *A. americana* assembled from genomic data differs by 2.2-13.5% from 13 sequences from *A. mustelae, A. marcianae, A. alata* and three unidentified species of *Alaria* (comparisons of various regions of the rDNA array, limited to sequences overlapping > 500 bp, AF184263, AY222091, JF769477-8, JF769480, JF769482, JF769484, JF820605, JF820607, JF820609, KT254014, KT254021, KT254023). Notably, *A. americana* differs from *A. marcianae* by 6.1% in partial 28S rDNA (914/973 identities with KT254021-2, Uhrig et al., 2015)) and by 8.4-8.9% in partial CO1 (KT254037-9, Uhrig et al. 2015).

#### 3.4.6 *Hysteromorpha triloba* (Rudolphi, 1819) (Fig. 8, Supplementary Fig. 3)

[Metacercaria; measurements from 7 paragenophores, ex lateral and cheek muscle of *Squalius cephalus* (mean weight 67 g) from Bidente River, Forlì-Cesena province, Emilia Romagna region, Italy; 10/10 fish infected with hundreds of metacercariae.]

Total length 776–889 (830±42, 7); body oval or pyriform, with poor demarcation between forebody and hindbody. Forebody round 536–664 (606±48, 7) long, 576–687 (630±33, 7) wide, with pseudosuckers forming cup-shaped depressions 48–80 (64±12, 7) deep, 44–96 (69±14, 12)wide. Hindbody bluntly triangular, roughly 120–303 (225±62, 7) long, 256–545 (404±96, 7) wide at widest point, where it joins forebody. Oral sucker terminal, 72–125 (82±19, 7) long, 52– 84 (70±12, 7) wide. Pharynx 50–76 (61±10, 7) long, 30–44 (36±4, 7) wide. Ventral sucker 60–82 (70±8, 7) long, 88–107 (99±6, 7) wide, sometimes anterior to, sometimes covered by tribocytic organ. Tribocytic organ trilobed, 160–229 (193±27, 6) long, 163–320 (207±54, 7) wide. Oesophagus 20–47 (34±19, 2) long, caeca almost reaching end of hindbody, flanking or passing ventrally over genital primordia in hindbody.

*Hysteromorpha corti* comb. nov. (Hughes, 1929) (Fig. 10)

**Fig 10.**
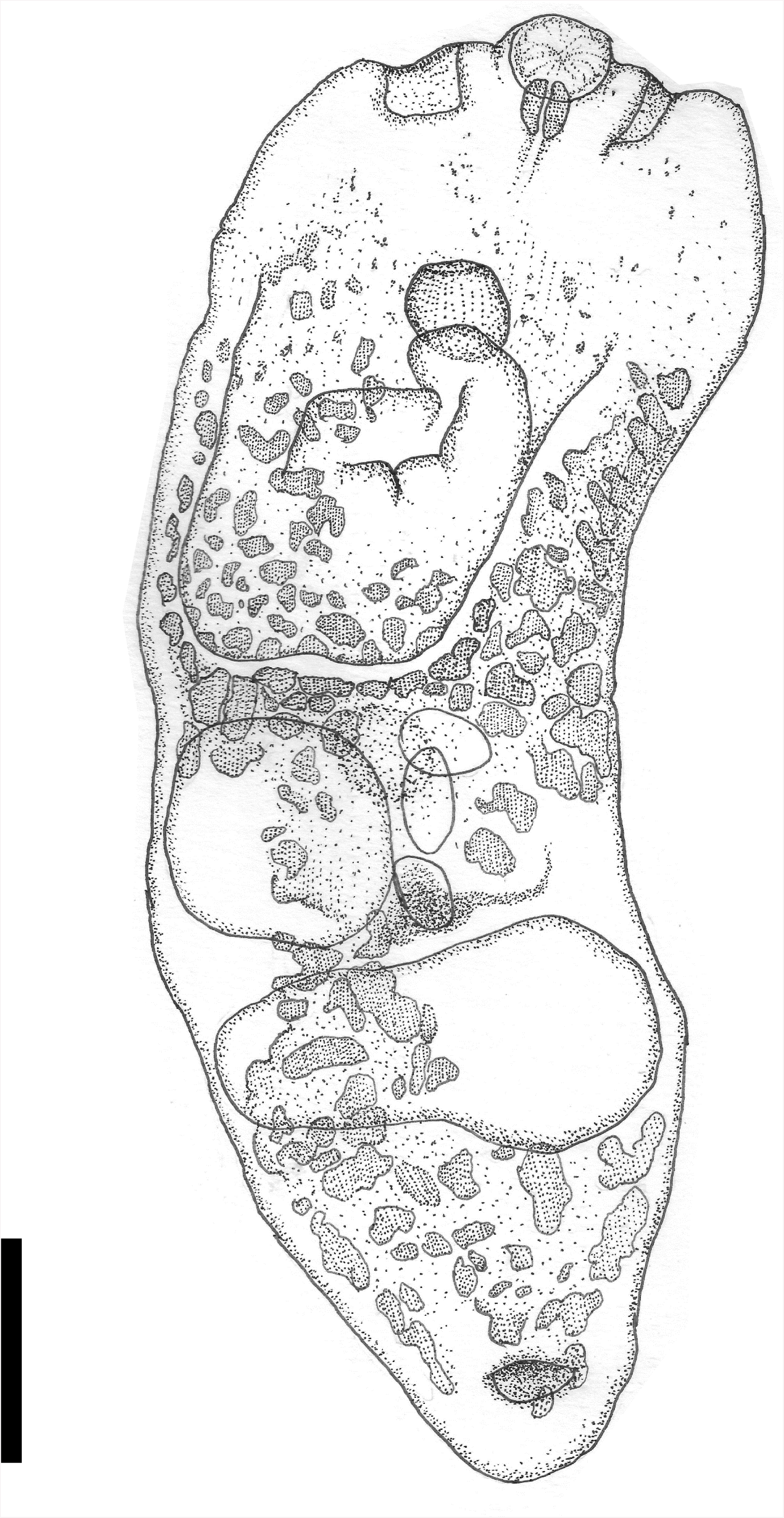
Adult of *Hysteromorpha corti* (Hughes, 1929) from *Phalacrocorax auritus*, Montreal, Quebec, Canada. Italy. Scale = 200 μm. Paragenophore for partial sequence of cytochrome *c* oxidase I, Genbank accession XZXXXXXX.

[Adult; measurements from 12 paragenophores, ex *Phalacrocorax auritus*, Montreal, Quebec, Canada]

Total length 1052 – 1633 (1314±172, 12); forebody and hindbody separated by constriction. Forebody spathulate, 490 - 762 (664±86, 12) long, 404 - 707 (509±82, 12) wide. Hindbody oval,490 - 943 (658±125, 12) long, 381 - 636 (480±68, 12) wide. Pseudosuckers forming recessed depressions in forebody, 47 - 129 (78±26, 9) deep, 70 - 200 (105±38, 9) across. Oral sucker 50 - 107 (75±16, 12) ×; 71 - 107 (86±12, 12). Ventral sucker 36 - 107 (75±18, 11) ×; 43 - 143 (93±30,11). Tribocytic organ 142 - 321 (241±58, 12) long, 190 - 293 (237±33, 11) wide. Pharynx muscular, pyriform, 52–68 (59±6, 5) long, 40–70 (52±11, 5) wide. Ovary 71 - 100 (85±14, 5) ×; 64 - 114 (81±19, 5), anterior to anterior testis. Anterior testis smooth, unevenly lobed, lateral, 107 - 214 (169±31, 10) ×; 107 - 250 (171±48, 7). Posterior testis extending laterally across hindbody width, with two lateral, round lobes, 143 - 229 (171±35, 9) ×; 357 - 500 (423±51, 8). Vitelline reservoir intertesticular; sub-median. Vitellaria from anterior to ventral sucker to posterior extremity, forming a narrow ventral band in hindbody at level of testes. Genital atrium subterminal, dorsal. Eggs 0–9, 87–109 (96±6, 12) long ×; 44–66 (54±6, 12) wide.

[Metacercaria; measurements from 7 syngenophores, ex *Catostomus commersoni* (n=1), *Notemigonus crysoleucas* (n=5), Montreal and Great Lakes region, Canada]

Total length 712–880 (796±55, 7); body oval or shield-shaped, with poor demarcation between forebody and hindbody. Forebody round 640–696 (670±25, 5) long, 384–472 (428±34, 7) wide. Hindbody bluntly triangular, roughly 80–160 (126±32, 5) long, 152–200 (176±34, 2) wide at widest point, where it joins forebody. Oral sucker terminal, 60–68 (62±4, 5) long, 56–68 (63±4,5) wide. Pharynx 32–35 (33±2, 3) in diameter. Ventral sucker 56–68 (60±6, 5) long, 60–76 (73±6, 6) wide, sometimes anterior to, sometimes partly covered by tribocytic organ. Tribocytic organ trilobed, 136–176 (163±18, 4) long, 116–160 (137±19, 5) wide.

##### 3.4.6.1 Diagnosis and Remarks

Sequences of CO1 from metacercariae of *H. triloba* from Italy differed by 6.9-9.7 (mean 8.7)% from material collected in North and Central America (Fig. 4). Among the Italian isolates, CO1 varied by 0.3-0.8 (mean 0.5)%, and among American samples, by 0-5.6 (mean 1.7)%. The rDNA operon of Italian *H. triloba* differed from American isolates by 0-0.2% from 18S and 28S subunit sequences 1281-1694 bp in length (HM114365, MF398354-7) and by 0.1-0.3% to ITS1- 5.8S-ITS2 sequences 1017-1261 in length (HM064925-7, JF769486, MG649479-93). These 20 aligned ITS sequences from *Hysteromorpha* had three variable sites, with one transitional mutation in ITS1 private to the Italian sequence. Rudolphi (1819) described *Hysteromorpha triloba* (as *Distoma trilobum*) in Europe from four adults from *Phalacrocorax carbo* collected by Bremser, who was based in Vienna (Sattmann et al., 2014). The type host has a wide, but mainly Eurasian distribution (Hatch et al., 2000). A description of metacercaria of *H. triloba* from cyprinids in Europe by Ciurea (1930) largely agrees with our observations of metacercariae from *S. cephalus* (Cyprinidae), and the ventral and oral suckers, pharynx and oesophagus are generally smaller in European than in American isolates (Supplementary Table 6). We consider these European and American isolates of *Hysteromorpha* to be separate species, even if meristic or morphological differences were not discerned between American and European adults of *Hysteromoprha* based on available data and descriptions (Supplementary Table 6), and rDNA divergence levels are small. Because the species was described in Europe, from a host with a Palaearctic distribution, and similar metacercariae are known from cyprinids in Europe (see also records in Bykhovskaya-Pavlovskaya, 1962), the name *H. triloba* is reserved for the Palearctic lineage. In the Nearctic, Hughes (1929) provided the first description of *Hysteromorpha*, from metacercariae in the muscle of *Ameiurus nebulosus* and *Ameiurus melas* in the Illinois River. Hughes (1929) named these metacercariae *Diplostomulum corti*. Hugghins (1954a, 1954b) synonymized *D. corti* with *Hysteromorpha triloba* (Rudolphi, 1819) Lutz 1931. The North American species of *Hysteromorpha* will therefore bear the name created by Hughes (1929).

#### 3.4.7 *Tylodelphys immer* Dubois, 1961 (Fig. 11)

[Measurements from 5 adult paragenophores, ex *Gavia immer*.]

**Fig 11.**
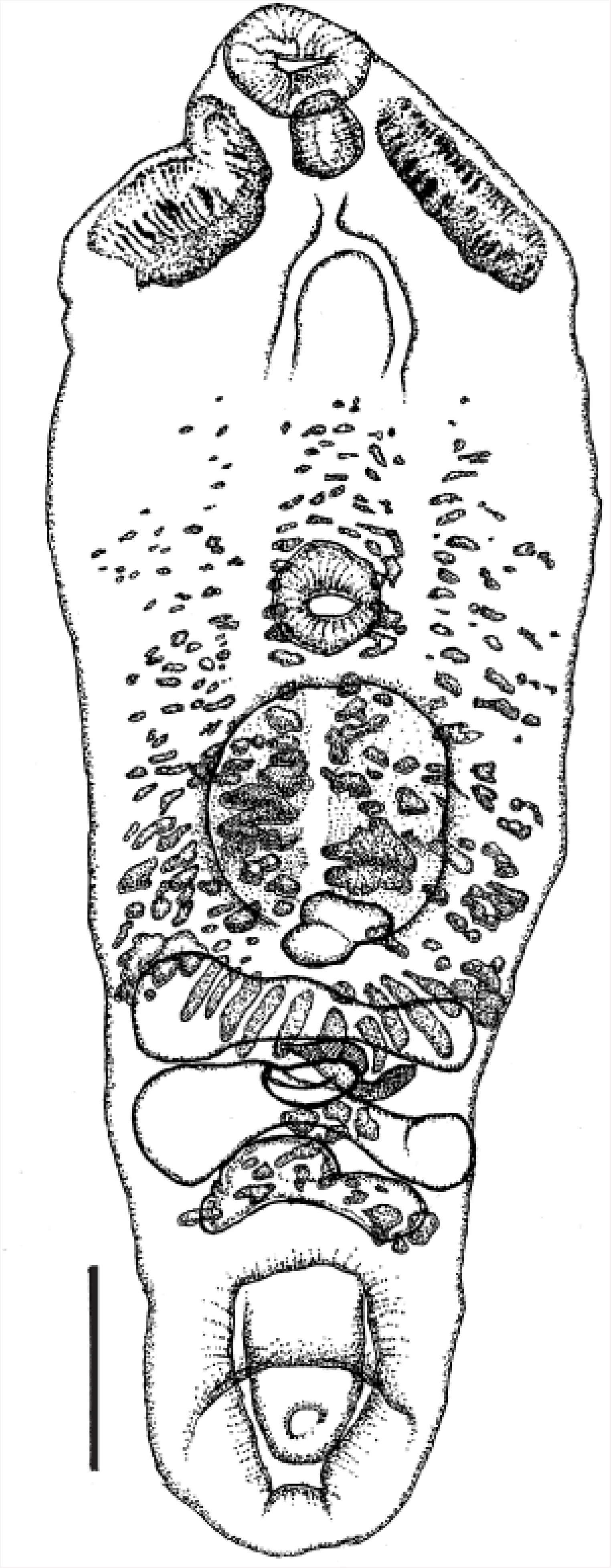
(a) Adult of *Tylodelphys immer* Dubois, 1961, from *Gavia immer* in Montreal, Quebec, Canada. Scale = 200 μm. (**b).** Ventral tegument of forebody. VS=ventral sucker, arrow points to anterior of worm. Scale = 50 μm. Paragenophore for partial sequence of cytochrome *c* oxidase I, Genbank accessions KR271483, KR271487, KR271489.

Total length 1515–1636; body linguiform, poor demarcation between forebody and hindbody. Forebody 970–1111 long, 455–556 wide, oval in shape with pointed anterior flanked by conspicuous pseudosuckers 160–232; forebody widest usually in anterior third (4/5 specimens). Ventral surface tegument with plicate folds on anterior half to two thirds of forebody. Cylindrical hindbody tapering to terminate in rounded extremity, 525–566 long, 284–404 wide. Hindbody length / forebody length 0.47–0.56; hindbody width / forebody width 0.51–0.85. Oral sucker terminal, 80–119 long, 100–127 wide. Pharynx well developed, 68–84 long, 66–80 wide. Ventral sucker 90–109 long, 100–131 wide, situated slightly more than halfway along length of forebody, the anterior edge occurring 51–54% from anterior end of forebody. Tribocytic organ oval to round, 240–288 long, 152–240 wide. Ovary round or bilobed, small, sub-median, near or overlapped by postero-lateral edge of tribocytic organ, 40–80 long, 56–80 wide. Testicular zone extending posteriorly from forebody-hindbody division, occupying first 29–39% of hindbody. Testes tandem, symmetric, bilobed. Anterior testis 80–96 long, 272–344 wide, wider than posterior testis. Posterior testis 88–112 long, 272–320 wide. Vitelline field densest in vicinity of tribocytic organ, extending two thirds of length of forebody, ending 31–34% of distance from anterior edge of ventral sucker to anterior extremity, in hindbody narrowing to a ventral strip between lobes of posterior testis, terminating in a lateral band at level of seminal vesicle. Vitelline reservoir intertesticular, sub-median. Copulatory bursa with subterminal, wide, thick-walled ventral opening 168–184 long, 160–184 wide, housing well-developed genital cone 112– 152 wide. Uterus with 1–8 eggs, 82–100 long ×; 44–68 wide.

#### 3.4.8 Remarks

Most of these observations and dimensions agree with the account of Dubois (1970b) (supplementary Table 7), although he did not comment on the tegument; a pseudosucker on one specimen was shorter (160 *versus* 180 minimum length in Dubois, 1970b); the pharynx was wider in three worms (74-80 *versus* maximum of 70 in Dubois, 1970b); the ventral sucker was longer in 4 worms (104–109 *versus* 100 in Dubois (1970b) and wider in one (131 *versus* 122 in Dubois, 1970b); the ovary was smaller (40–80×56–80 *versus* 95–115×80–145 in Dubois, 1970b) and more median in 3 worms; the anterior testis was shorter in 3 worms (80-88 versus 90 in Dubois, 1970b), the posterior in one (88 versus 90 in Dubois, 1970b); and the genital cone was wider in 2 worms (152 *versus* 135 in Dubois, 1970b). These variations seem within that to be expected within species, and thus not taxonomically significant. They could reflect differences in maturity, as the worms included in our analysis had 0-8 eggs, while Dubois (1970b) examined material with 4-17 eggs, or again might be artefacts of differences in specimen preparation.

The specimens used for morphological and genomic analysis in the present study were from the same individual host as the *T. immer* studied by Locke et al. (2015), and the mitochondrial genome and rDNA operon were 98.1-99.9 % similar to CO1 (Fig. 4) and identical to ITS sequences (KT186804-6) of Locke et al. (2015).

## 4. Discussion

We confirmed and expanded on recent analyses showing a paraphyletic pattern of mt genome evolution in the Diplostomida (Brabec et al., 2015; Briscoe et al., 2016; Chen et al., 2016). The mt genomic phylogeny conflicts with the rDNA phylogeny upon which the Diplostomida was erected (Figs. 1, 2). There were about one quarter as many variable sites in the rDNA alignment as in our alignment of mt nucleotides (1444 variable sites in Olson et al., 2003). Thus, the well-supported mt topology in Fig. 2, containing members of two of three superfamilies in the Diplostomida, might have cast doubt on the validity of the order. However, a much larger genomic dataset, which we had designated as an arbiter between mitonuclear alternatives, yielded unequivocal support for the order (Fig. 3).

Discordance between nuclear and mt phylogenies is not uncommon, and although it is more often recorded at shallower nodes than in the present study (e.g., Perea et al., 2016; Platt et al., 2018), differences also occur among deeply divergent lineages (e.g., Sun et al., 2015, and compare Inoue et al., 2003 and Faircloth et al., 2013). In the present case, the discrepancy occurs along short internal branches at the base of longer terminal branches (Figs. 2, 3). This is consistent with ancient, rapid radiation, which is inherently difficult to resolve, particularly in conjunction with incomplete lineage sorting (Whitfield and Lockhart, 2007). Along these short internal branches, mitochondrial genomes of digeneans may have a lower phylogenetic signal/noise ratio than nuclear genes, exacerbating effects of incomplete taxon sampling (Graybeal and Cannatella, 1998; Hedtke et al., 2006; Philippe et al., 2011). Genomic data (both mt and nuclear) from other diplostomidans and early, divergent lineages from the Plagiorchiida are needed to clarify this (Brachylaimoidea Joyeux and Foley, 1930, Liolopidae Odhner, 1912, Aporocotylidae Odhner, 1912, Spirorchiidae Stunkard, 1921, Bivesiculoidea Yamaguti, 1934).

Within the Diplostomoidea, however, the mt and UCE phylogenies are congruent, which suggests they reflect evolutionary relationships among the species studied. Figures 2 and 3 also share elements that recur in prior work. In most molecular phylogenies, the superfamily is monophyletic and cyathocotylids are basal (Blasco-Costa and Locke, 2017; Dzikowski et al. 2004; Hernández-Mena et al., 2017), as herein. Both the branching order and composition of the major diplostomoid clades in Figs. 2 and 3 herein are consistent with an analysis of concatenated CO1 and rDNA spacer sequences by Blasco-Costa and Locke (2017). The association of *Hysteromorpha, Alaria, Tylodelphys* and *Diplostomum* is consistent with trees in Fraija-Fernández et al. (2015), López-Jiménez et al. (2017), and Olson et al. (2003). Discrepancies (e.g., *Hysteromorpha* in Hernández-Mena et al., 2017) are typically associated with poor nodal support. These genera are also consistently associated with *Austrodiplostomum* and *Neodiplostomum* (Blasco-Costa and Locke, 2017; Hernández-Mena et al., 2017; Locke et al., 2015; the latter sometimes misidentified as *Fibricola*, see Blasco-Costa and Locke, 2017), and the name Diplostomidae should be reserved for members of this group.

Like other studies, our analysis indicates the Diplostomidae *s.l.* is paraphyletic. *Posthodiplostomum*, though nominally part of the family, is separate from and basal to a clade composed of other diplostomids and the Strigeidae. *Posthodiplostomum* belongs to the Crassiphialinae Sudarikov, 1960 and is consistently recovered with other members of this subfamily (e.g., *Uvulifer, Bolbophorus, Ornithodiplostomum, Mesoophorodiplostomum* and *Posthodiplostomum* in Athokpam and Tandon, 2014; Blasco-Costa and Locke, 2017; Hernández-Mena et al., 2017; López-Jiménez et al., 2017; Sereno-Uribe et al., 2018; Locke et al., 2010, see fig. S4). One recent analysis suggests some of these genera are not distinct (López-Hernández et al., 2018). As noted by Blasco-Costa and Locke (2017) and Hernández-Mena et al. (2017), it appears that the Crassiphialinae will rise to the family level, although it would be prudent to obtain data from the type genus, *Crassiphiala* Van Haitsma, 1925, before enacting this.

Both the present and the weight of prior phylogenetic analysis (see above) suggest that some family-level clades in the Diplostomoidea are distinguishable by long-recognized metacercarial morphotypes. For example, crassiphialinids and other diplostomids are clearly evolutionarily distinct, and readily separated by their metacercariae (neascus and diplostomulum). In adult forms, members of these two clades are sometimes discriminated by the distribution of the vitellaria, but this character fails in many cases (Niewiadomska, 2002c). The view that the metacercaria offers an impoverished subset of the phylogenetically informative characters found in the adult (Gibson, 1987) seems not to apply in the Diplostomoidea. For example, the infection sites and encystment habits of metacercariae may be phylogenetically conserved. Diplostomid genera with metacercariae that reside unencysted in the eyes of second intermediate hosts consistently group together in molecular phylogenies (Blasco-Costa and Locke, 2017; Hernández-Mena et al., 2017) and in *Diplostomum*, habitats within the eye are conserved (Blasco-Costa et al., 2014) and may influence diversification (Locke et al., 2015). Some morphological characters are also more easily visualized in metacercariae. For example, the character that mapped best onto phylogenetic analysis herein, the structure of the reserve bladder, is seldom described in the adult, likely because it is obscured by reproductive structures in mature worms (for an exception, see Overstreet et al., 2002). The value of this character fits Niewiadomska’s (2002a) concept of four main morphotypes. Shoop (1989) distinguished additional types of metacercariae, but molecular phylogenies have not supported their distinctness (e.g., see *Neodiplostomum/Fibricola*, and *Bolbophorus*, which possess neo- and prodiplostomula in Shoop’s system, in Blasco-Costa and Locke, 2017; Hernández-Mena et al., 2017). Further assessing the evolution of metacercarial characters and morphotypes will require additional molecular and morphological analysis. In the Strigeidae, all metacercariae are of a single type (tetracotyle), but the family is frequently non-monophyletic in molecular studies (Blasco-Costa and Locke, 2017; Hernández-Mena et al., 2017). It would be fruitful to characterize the reserve bladder in metacercariae belonging to the Proterodiplostomidae, which Hernández-Mena et al. (2017) found to be an early diverging, but not basal member of the diplostomoid clade. The simpler reserve bladder in the Cyathocotylidae, and the reticulate forms with transverse commisures in the crown clade of Strigeidae and Diplostomidae, predict intermediate complexity in proterodiplostomids.

The initial aim of this study was a phylogenomic evaluation of higher relationships among a small number of worms. To this end, we selected seven specimens identified to genus that were promising for shotgun sequencing. After closer examination of vouchers, we recorded a new species of *Cotylurus*, resurrected a species of *Hysteromorpha*, and found support for a species of *Alaria* of contested validity, and these findings were supported with molecular data. Identifications were less than straightforward in 3/7 cases, a proportion similar to the 20/44 studies in which diplostomoid diversity differed from expectations (reviewed by Blasco-Costa and Locke, 2017). One taxonomic result involved reconsideration of *H. triloba*, which has long been thought to be cosmopolitan (Hugghins, 1954b; Locke et al., 2011; Lutz, 1931; Sereno-Uribe et al., 2018). The genetic divergence seen in *Hysteromorpha* could be construed as intraspecific variation in widely separated populations (particularly the low level of rDNA variation), but we believe recently formed species to be a more plausible explanation. In addition to the molecular evidence, the non-overlapping ranges of the definitive hosts of Nearctic and Paleartic *Hysteromorpha* (i.e., *Phalacrocorax* spp.) suggest long isolation, and the distinction between *H. triloba* and *H. corti* are supported by morphological differences in the metacercaria. Moreover, the finding that *Hysteromorpha* is represented by a distinct species in North America, *H. corti*, is consistent with a general trend. With sequences now available from thousands of specimens and many valid and putative species of diplostomoids and clinostomids, intercontinental distributions supported by molecular data are exceedingly rare, and limited to distributions along the margins of a second continent (e.g., *Austrodiplostomum compactum* (=*A. ostrowskiae*), Locke et al. 2015). More commonly, a single species thought to cosmopolitan is revealed by DNA to be comprised of multiple geographically isolated species (e.g., *Diplostomum spathaceum, D. baeri, Clinostomum complanatum*, Caffara et al., 2011; Galazzo et al., 2002; Locke et al., 2015). The presence of the North American species, *P. centrarchi*, in Europe (Kvach et al., 2017; Stoyanov et al., 2017) and the Caribbean (Bunkley-Williams and Williams, 1994; present study) is instructive, as it is associated with recent introductions of non-native intermediate hosts. The overall pattern suggests intercontinental distributions based on historical records should be regarded skeptically in diplostomoids, clinostomids, and, we would suggest, other digeneans with life cycles tied to fresh water. For example, we predict that DNA will reveal that Palearctic *Apharyngostrigea cornu* is distinct from North American isolates sequenced by Locke et al. (2011). Because *A. cornu* was described in Europe (Zeder, 1800), the North American lineage will need to be renamed. Similarly, the Holarctic distribution of *Cotylurus brevis* (Dubois, 1970b; McDonald, 1981) seems doubtful. South American isolates of *H. triloba* (Lunaschi et al., 2007; Lutz, 1931) are also likely to represent another species, distinct from the North American and European lineages.

## Acknowledgements

We are indebted to Brandon Ballengée (McGill University), Kimberly Bates (Winona State University), Matías J. Cafaro (University of Puerto Rico at Mayagüez, UPRM), Gary Conboy (University of Prince Edward Island), Martin Kalbe (Max Planck Institute for Evolutionary Biology), David Kerstetter (Nova Southeastern University), Audrey J. Majeske (UPRM), J. Daniel McLaughlin (Concordia University), and Le Nichoir Wildlife Rehabilitation Centre for providing hosts, worms, or laboratory access. Lutz Froenicke and staff at the UC Davis Genome Center are gratefully acknowledged. Isabel Blasco-Costa (Natural History Museum of Geneva) provided constructive suggestions that improved an earlier version of the manuscript. Early phases of this study were supported by the Canadian Federal Government’s Genomics Research Development Initiative, an NSERC Discovery Grant (A6979), and by Paul D. N. Hebert at the Center for Biodiversity Genomics, University of Guelph, Canada through funding from NSERC, Genome Canada, the Ontario Genomics Institute and the International Barcode of Life initiative. Major funding support was provided by the Puerto Rico Science, Technology and Research Trust to SL and computational support was provided to by an NSF-XSEDE grant (TG-BIO170048) SL and AVD.

## Figure legends

**Supplementary Fig 1:**
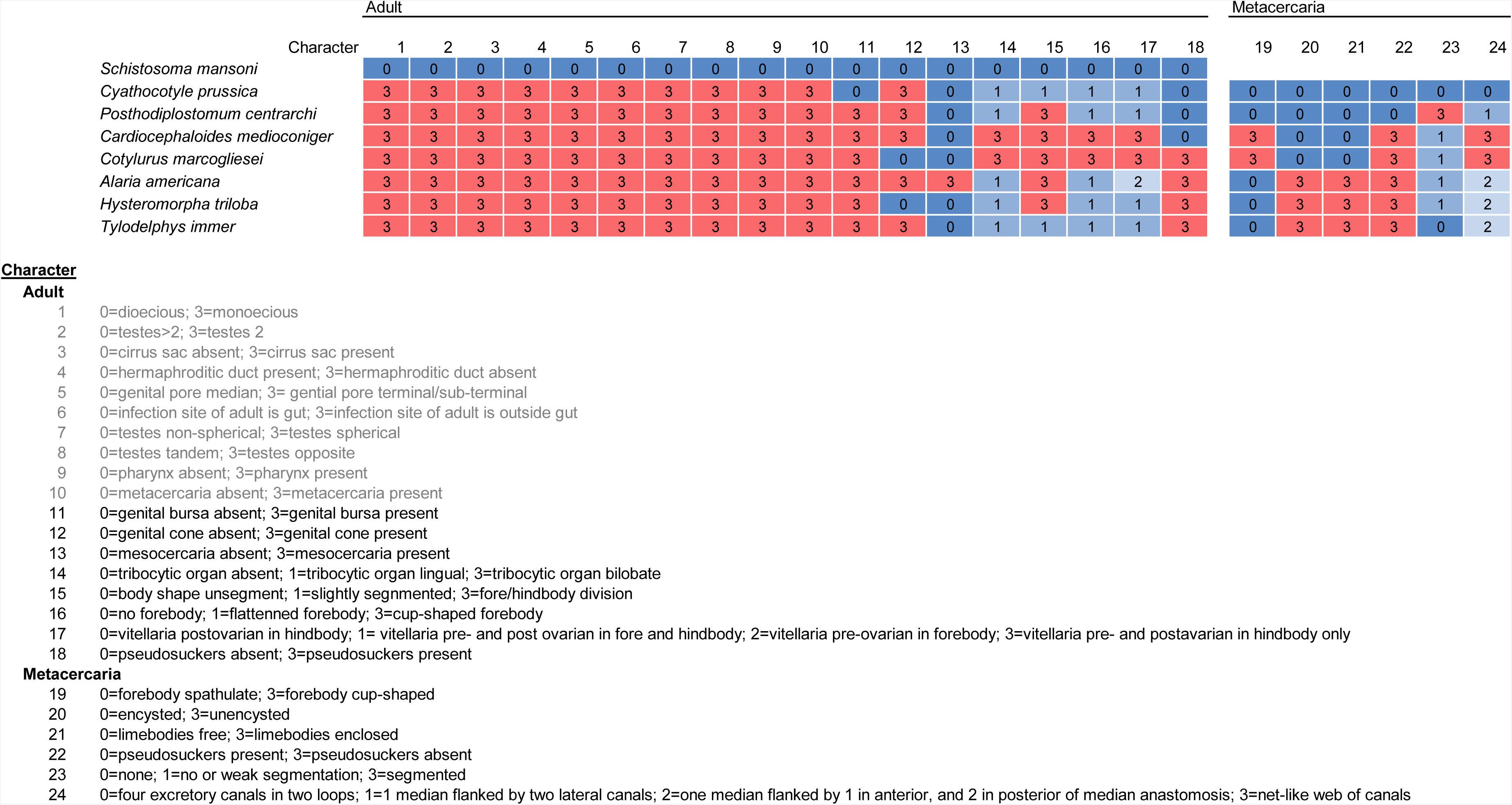
Characters mapped onto the topology of the phylogenomic analysis of the Diplostomoidea.

**Supplementary Fig 2:**
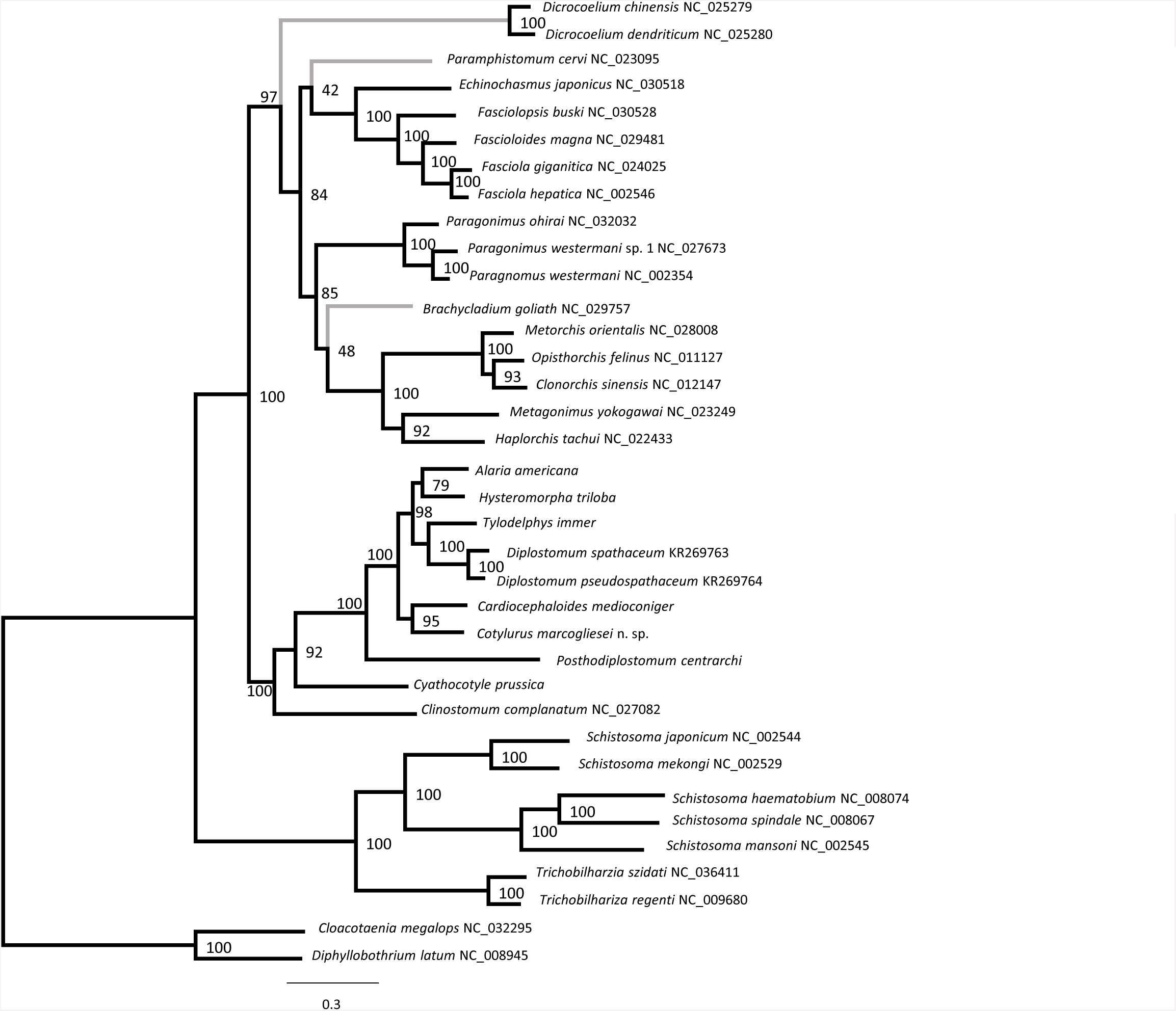
Phylogenetic analysis of seven representatives of the Diplostomoidea and 29 other members of the Platyhelminthes, estimated using maximum likelihood based on translated amino acids in 13 protein-coding genes in the mitochondrion

**Supplementary Fig 3:**
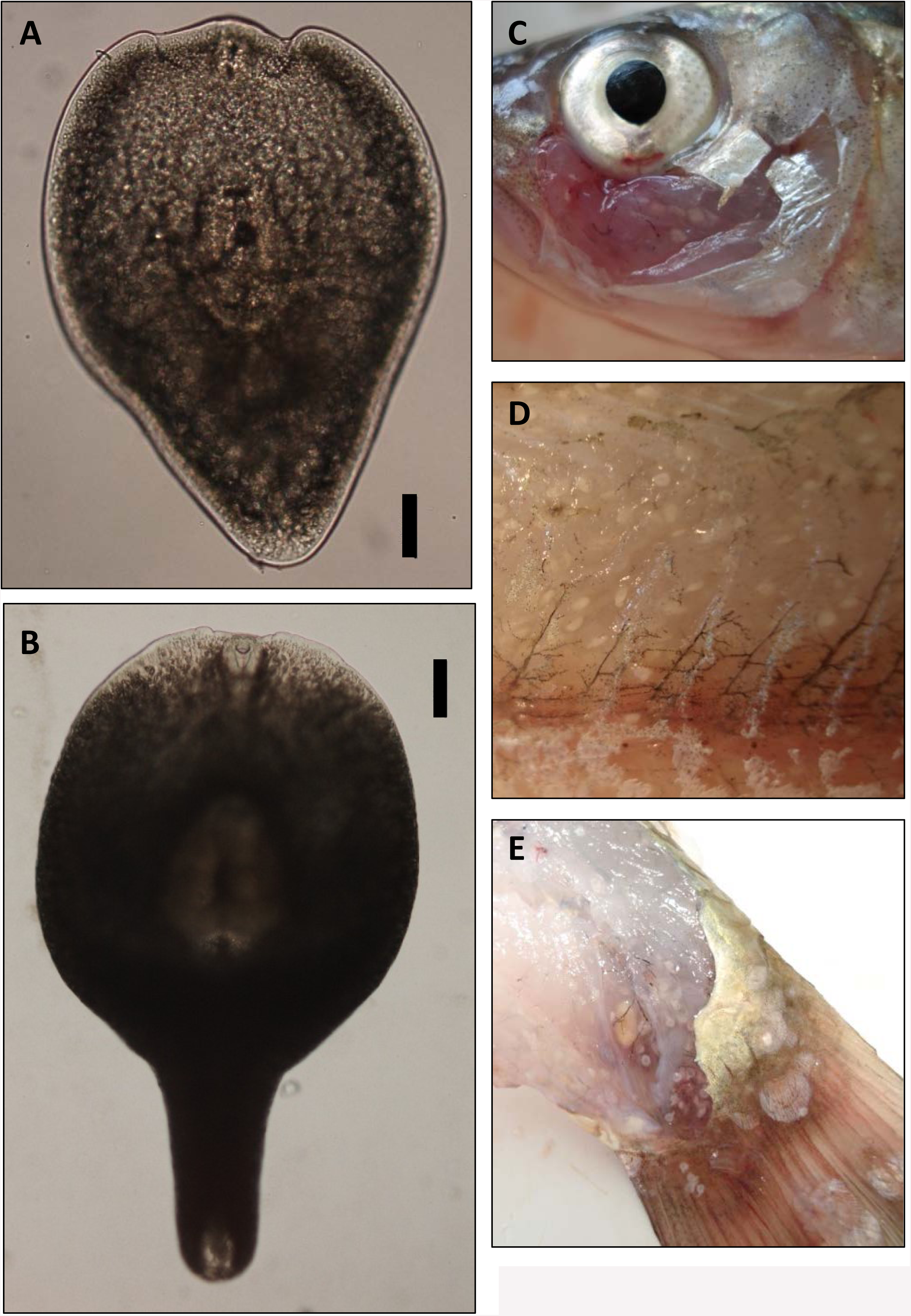
Metacercaraie of *Hysteromorpha triloba* in muscle of *Squalius cephalus*. A-B: live metacercariae (scale = 100 μm); Metacercariae in cheek (C), lateral (D) and caudal (E) muscle.

---

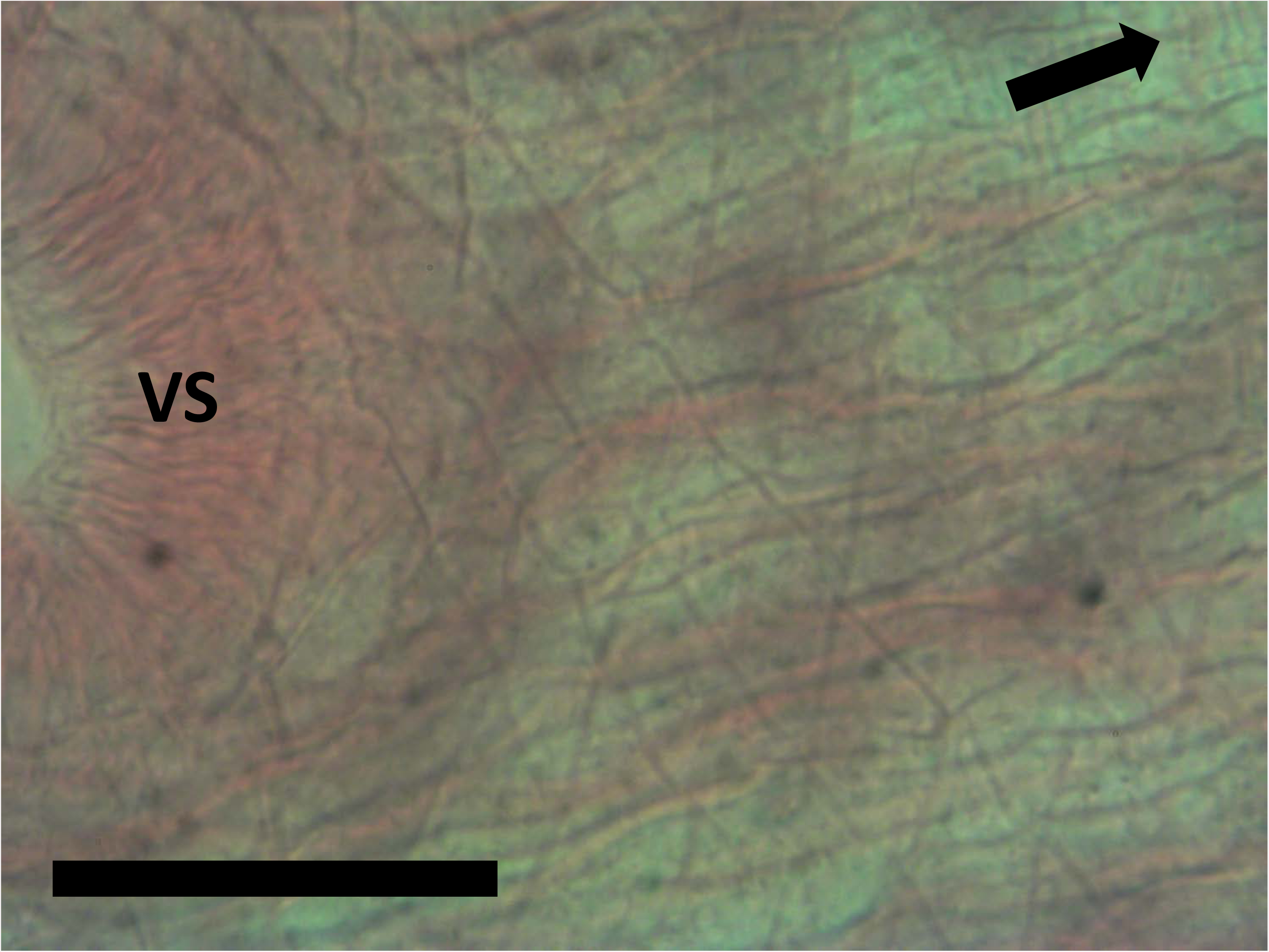

**Supplementary Table 1:**
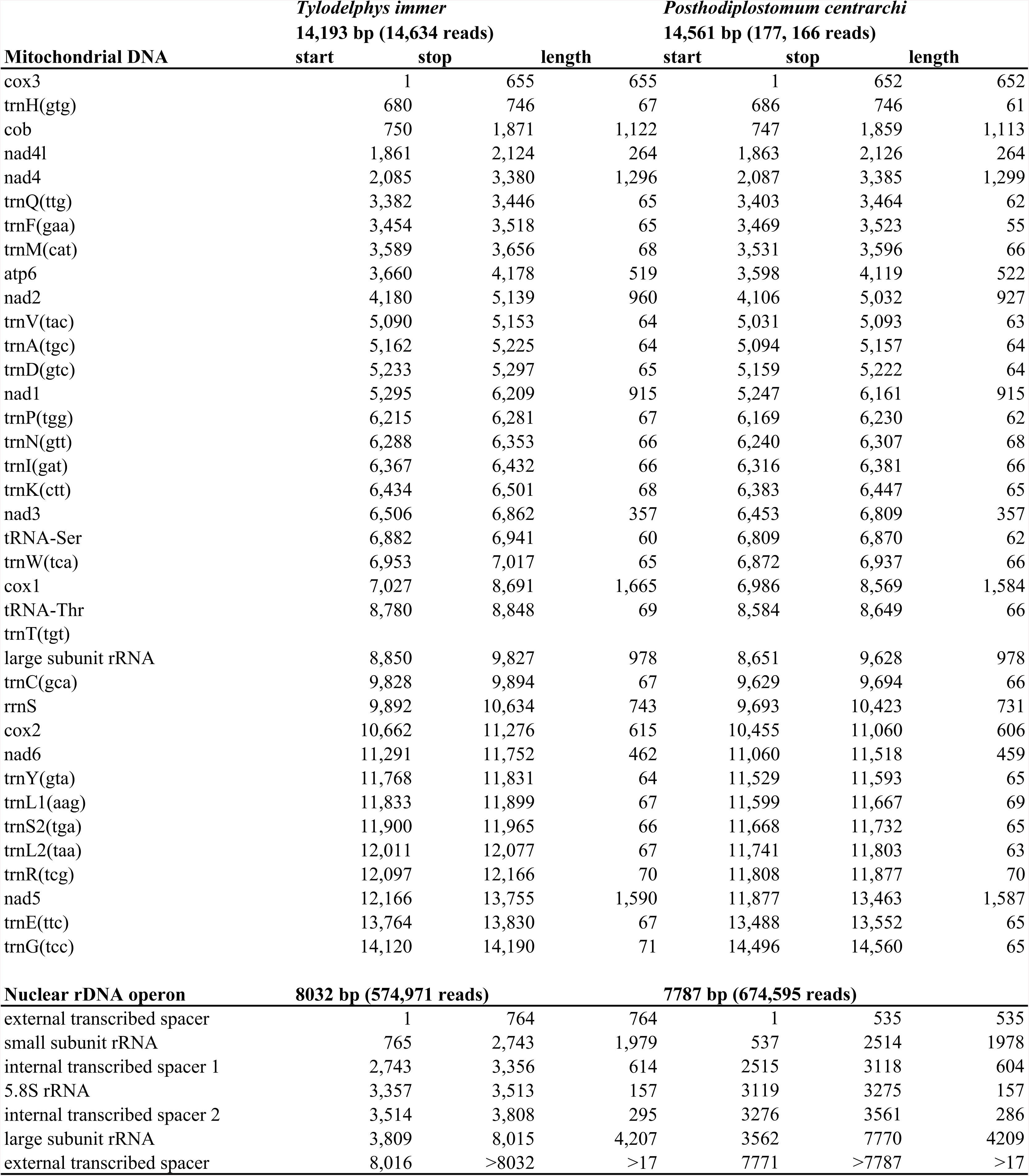
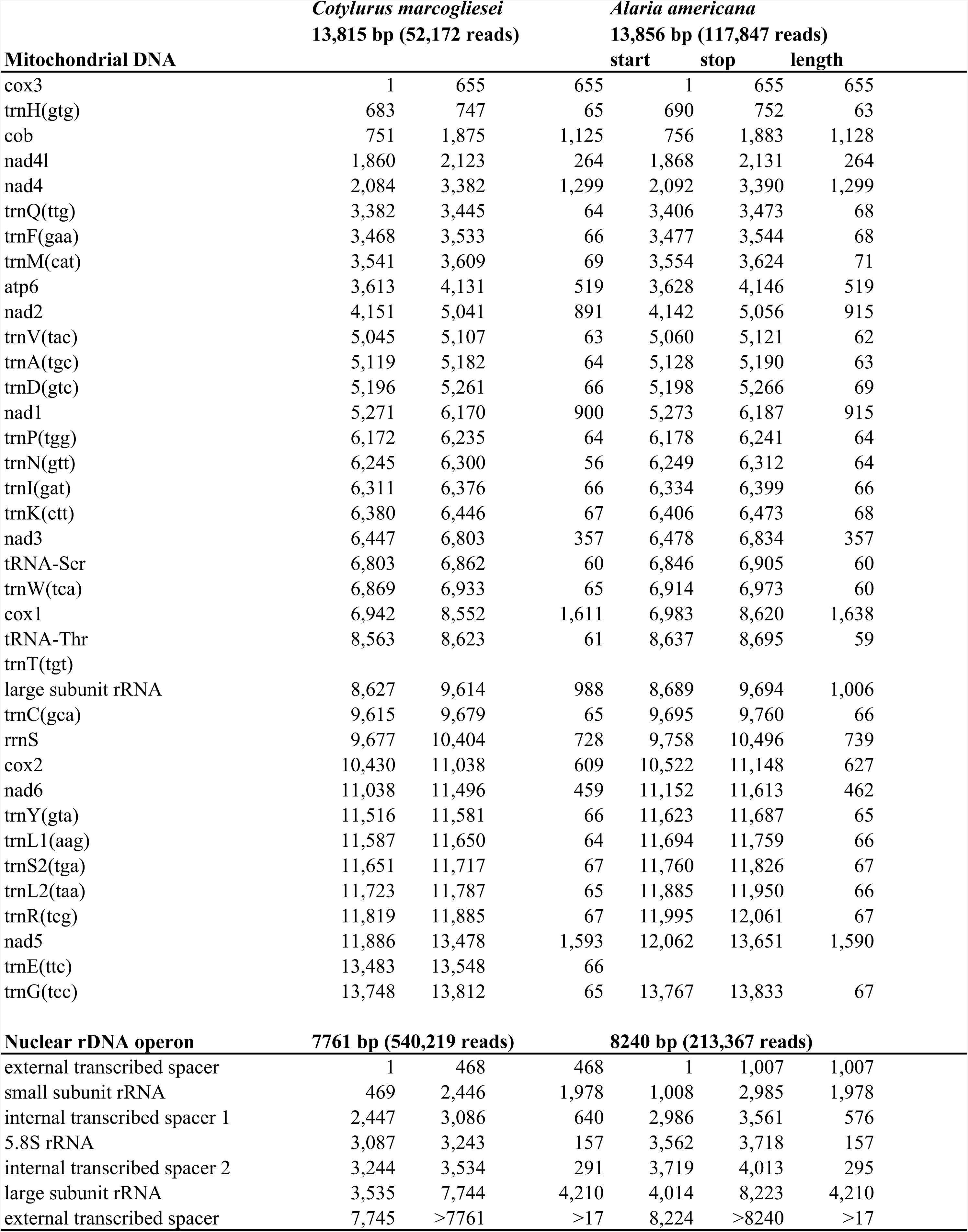
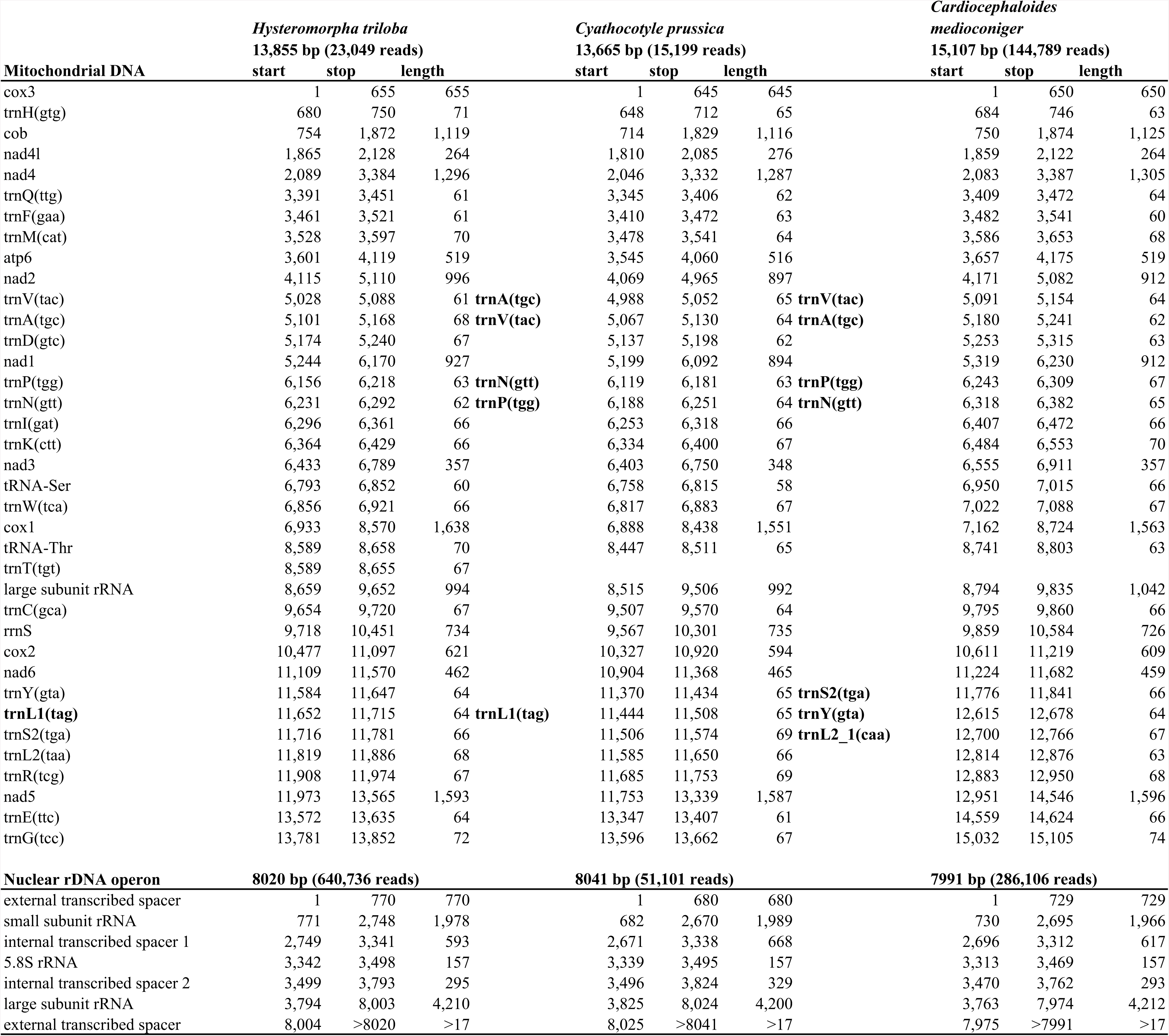
Characteristics of mitochondrial genomes and rDNA operons for seven members of the Diplostomoidea.

**Supplementary Table 2:** Selected morphometrics from adults of *Posthodiplostomum* reported in μm as range (mean, ± standard deviation, n).

| Identification (present study) | *Posthodiplostomum centrarchi* |  | *P. centrarchi* and/or<br>*Posthodiplostomum minimum* |  |
| --- | --- | --- | --- | --- |
| Identification (original source) |  |  | *P. minimum centrarchi*, and/or<br>*P. minimum minimum* |  |
| Source | present study |  | Dubois, 1968 |  |
| Life stage | adult |  | adult |  |
| Host | *Ardea herodias* |  |  |  |
| Locality | Montreal, Quebec, Canada |  |  |  |
|  | Length | Width | Length | Width |
| Body | 1222 - 1775 (1518, ±186, 9) | | 890 - 1700 | |
| Forebody | 680 - 1200 (911, ±165, 9) | 452 - 875 (622, ±130, 9) | 540 - 1150 | 250 - 600 |
| Hindbody | 520 - 875 (639, ±108, 9) | 248 - 750 (373, ±159, 9) | 300 - 600 | 160 - 470 |
| Forebody/Hindbody | 0.58 - 0.86 (0.71, ±0.09, 9) | | 0.41 - 0.83 | |
| Oral sucker | 38 - 72 (52, ±10, 8) | 38 - 72 (56, ±10, 8) | 30 - 66 | 30 - 60 |
| Pharynx | 40 - 53 (49, ±4, 8) | 40 - 53 (42, ±5, 8) | 26 - 53 | 24 - 47 |
| Oesophagus | 60 - 72 (66, ±8, 2) | | 25 - 90 | |
| Ventral sucker | 55 - 90 (76, ±11, 9) | 55 - 80 (71, ±8, 7) | 42 - 95 | 50 - 100 |
| VS position % in forebody | 54 - 71 (60, ±6, 7) | | 60 - 71 | |
| Tribocytic organ | 140 - 200 (178, ±22, 8) | 144 - 256 (195, ±44, 6) | 125 - 220 | 125 - 190 |
| Ovary | 80 - 105 (92, ±11, 4) | 72 - 88 (80, ±6, 4) | 35 - 100 | 42 - 116 |
| Anterior testis | 96 - 200 (159, ±33, 7) | 144 - 272 (204, ±49, 7) | 70 - 170 | 120 - 240 |
| Posterior testis | 176 - 350 (257, ±58, 8) | 216 - 350 (280, ±43, 9) | 70-240 | 170-330 |
| Eggs | 70 - 98 (84, ±11, 10) | 42 - 64 (56, ±8, 10) | 73 - 91 | 48 - 64 |
| N eggs | 0 - 4 (1, ±1.7, 9) | | ≤8 | |
| Copulatory bursa | 120 - 216 (177, ±37, 8) | 135 - 300 (248, ±55, 9) | 145 - 160 | |

**Supplementary Table 3:**
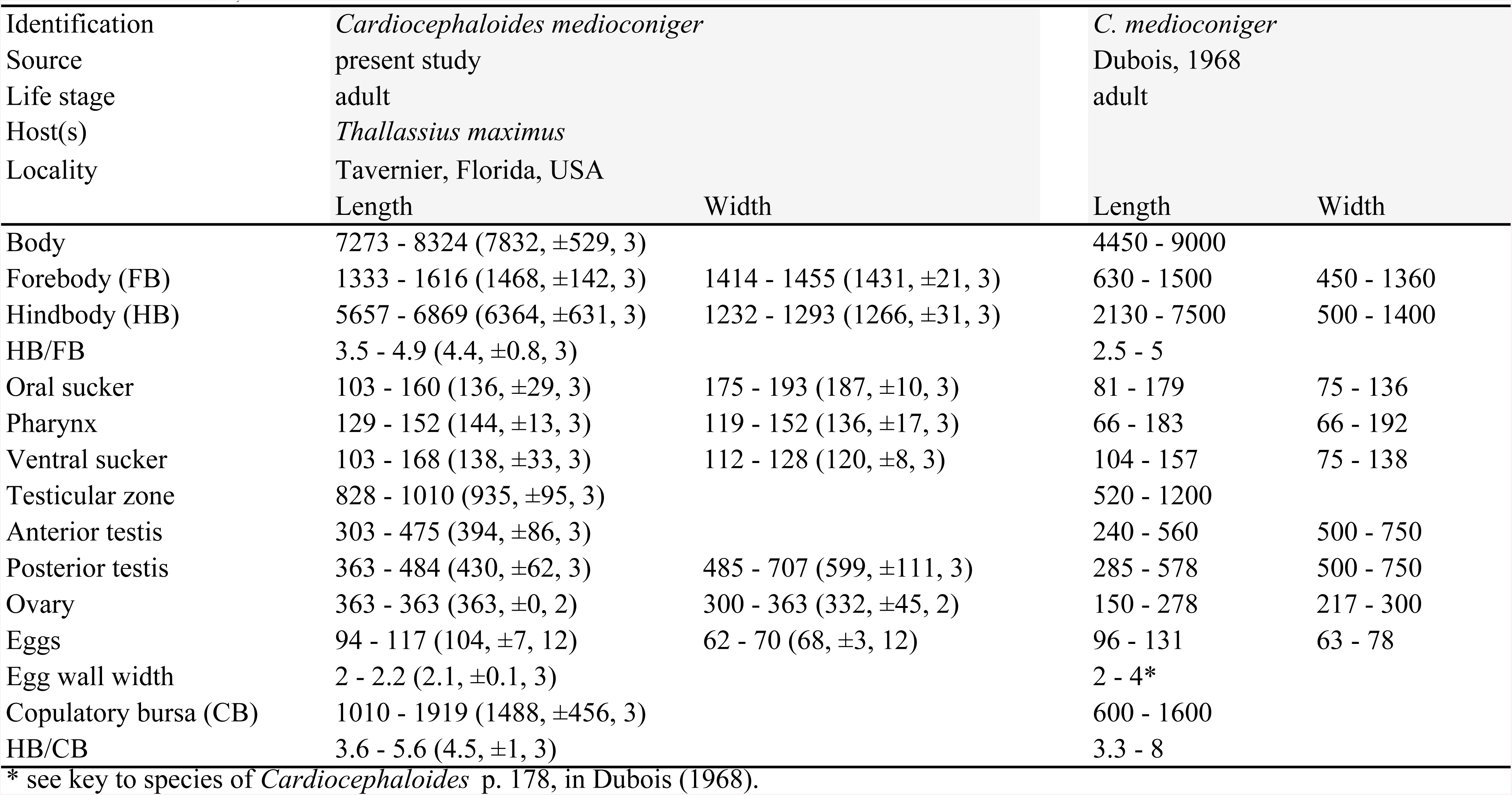
Selected morphometrics from adults of *Cardiocephaloides medioconiger* reported in μm as range (mean, ± standard deviation, n).

| Identification | *Cardiocephaloides medioconiger* |  | *C. medioconiger* |  |
| --- | --- | --- | --- | --- |
| Source | present study |  | Dubois, 1968 |  |
| Life stage | adult |  | adult |  |
| Host(s) | *Thalassius maximus* |  |  |  |
| Locality | Tavernier, Florida, USA |  |  |  |
|  | Length | Width | Length | Width |
| Body | 7273 - 8324 (7832, $\pm 529$, 3) | | 4450 - 9000 | |
| Forebody (FB) | 1333 - 1616 (1468, $\pm 142$, 3) | 1414 - 1455 (1431, $\pm 21$, 3) | 630 - 1500 | 450 - 1360 |
| Hindbody (HB) | 5657 - 6869 (6364, $\pm 631$, 3) | 1232 - 1293 (1266, $\pm 31$, 3) | 2130 - 7500 | 500 - 1400 |
| HB/FB | 3.5 - 4.9 (4.4, $\pm 0.8$, 3) | | 2.5 - 5 | |
| Oral sucker | 103 - 160 (136, $\pm 29$, 3) | 175 - 193 (187, $\pm 10$, 3) | 81 - 179 | 75 - 136 |
| Pharynx | 129 - 152 (144, $\pm 13$, 3) | 119 - 152 (136, $\pm 17$, 3) | 66 - 183 | 66 - 192 |
| Ventral sucker | 103 - 168 (138, $\pm 33$, 3) | 112 - 128 (120, $\pm 8$, 3) | 104 - 157 | 75 - 138 |
| Testicular zone | 828 - 1010 (935, $\pm 95$, 3) | | 520 - 1200 | |
| Anterior testis | 303 - 475 (394, $\pm 86$, 3) | | 240 - 560 | 500 - 750 |
| Posterior testis | 363 - 484 (430, $\pm 62$, 3) | 485 - 707 (599, $\pm 111$, 3) | 285 - 578 | 500 - 750 |
| Ovary | 363 - 363 (363, $\pm 0$, 2) | 300 - 363 (332, $\pm 45$, 2) | 150 - 278 | 217 - 300 |
| Eggs | 94 - 117 (104, $\pm 7$, 12) | 62 - 70 (68, $\pm 3$, 12) | 96 - 131 | 63 - 78 |
| Egg wall width | 2 - 2.2 (2.1, $\pm 0.1$, 3) | | 2 - 4* | |
| Copulatory bursa (CB) | 1010 - 1919 (1488, $\pm 456$, 3) | | 600 - 1600 | |
| HB/CB | 3.6 - 5.6 (4.5, $\pm 1$, 3) | | 3.3 - 8 | |

\* see key to species of *Cardiocephaloides* p. 178, in Dubois (1968).

**Supplementary Table 4:**
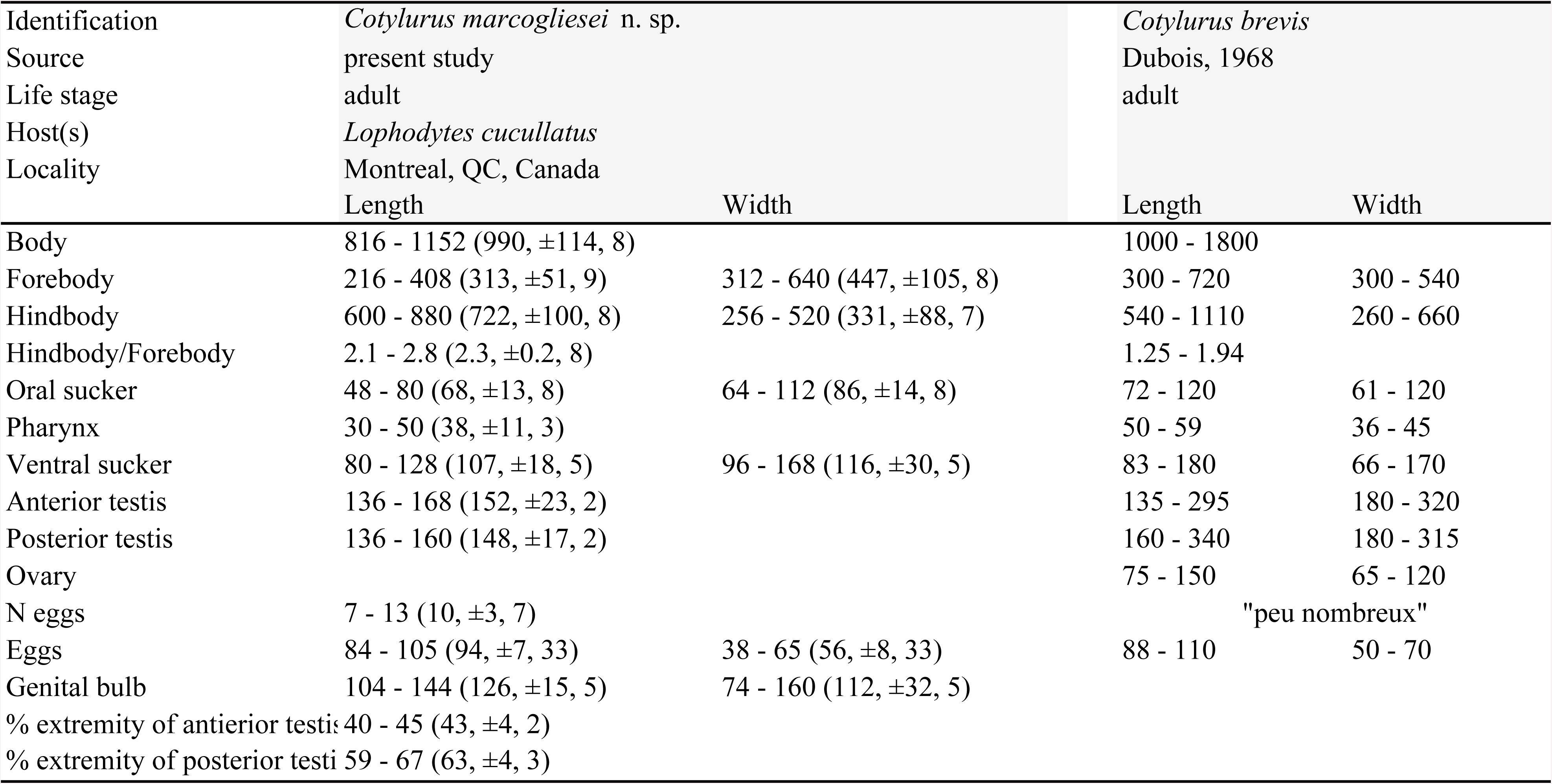
Selected morphometrics from adults of *Cotylurus* reported in μm as range (mean, ± standard deviation, n).

**Supplementary Table 5:** Selected morphometrics from adults of *Alaria* reported in μm as range (mean, ± standard deviation, n).

| Identification | *Alaria americana* |  | *Alaria americana* |  | *Alaria americana* (= *Alaria canis*) |  |
| --- | --- | --- | --- | --- | --- | --- |
| Source | present study |  | sources cited in Johnson, 1968 |  | Larue and Fallis, 1936 |  |
| Life stage | adult |  | adult |  | adult |  |
| Host(s) | *Vulpes vulpes* |  | *Canis familiaris*, *Vulpes fulva* |  | *Canis familiaris* |  |
| Locality | Montreal, QC, Canada |  |  |  |  |  |
|  | Length | Width | Length | Width | Length | Width |
| Body | 2121 - 2868 (2337, $\pm 275$, 7) | | 1160 - 4200 | | 2500 - 4200 (3200) | |
| Forebody (FB) | 1375 - 1858 (1510, $\pm 177$, 7) | 465 - 869 (592, $\pm 153$, 6) | | | 1600 - 2600 (2200) | 680-950 (800) |
| Hindbody (HB) | 606 - 1010 (815, $\pm 124$, 8) | 252 - 559 (458, $\pm 99$, 7) | | | 680 - 1600 (1000) | 750 - 1100 (920) |
| FB/HB | 1.8 - 2.3 (1.9, $\pm 0.24$, 7) | | 1.4 - 2.2 | | | |
| Lappet | 136 - 240 (190, $\pm 38$, 5) | | | | | |
| Oral sucker (OS) | 64 - 120 (89, $\pm 20$, 5) | 50 - 119 (82, $\pm 27$, 5) | 75 - 140 | 75 - 150 | 75 - 140 (100) | 110 - 150 (120) |
| Pharynx (PH) | 120 - 150 (134, $\pm 11$, 6) | 45 - 96 (81, $\pm 20$, 6) | 120 - 196 | 80 - 153 | 140 - 170 (150) | 90 - 140 (120) |
| OS/PH | 0.49 - 0.83 (0.66, $\pm 0.14$, 5) | | 0.67 | | | |
| Ventral sucker | 88 - 104 (95, $\pm 6$, 6) | 88 - 120 (101, $\pm 11$, 6) | 70 - 176 | 90 - 140 | 90 - 120 (110) | 90 - 140 (110) |
| Tribocytic organ | 667 - 788 (736, $\pm 40$, 6) | 160 - 250 (218, $\pm 40$, 4) | | | | |
| Ovary | 67 - 160 (105, $\pm 37$, 5) | 65 - 280 (172, $\pm 92$, 4) | | | | |
| Anterior testis | 110 - 319 (219, $\pm 91$, 4) | 120 - 327 (228, $\pm 104$, 3) | | | | |
| Posterior testis | 80 - 283 (165, $\pm 79$, 5) | 152 - 270 (228, $\pm 66$, 3) | | | | |
| Ejaculatory pouch | 250 - 444 (346, $\pm 70$, 8) | 101 - 135 (119, $\pm 10$, 8) | | | | |
| Ejaculatory pouch wall | | 31 - 55 (42, $\pm 7$, 8) | | 35 - 55 | | |
| Eggs | 102 - 136 (117, $\pm 8$, 25) | 36 - 80 (64, $\pm 8$, 25) | 90 - 133 | 64 - 86 | 108 - 116 (113) | 64 - 76 (70) |
| N eggs | 0 - 11 (7, $\pm 4$, 8) | | | | | |

**Supplementary Table 6:**
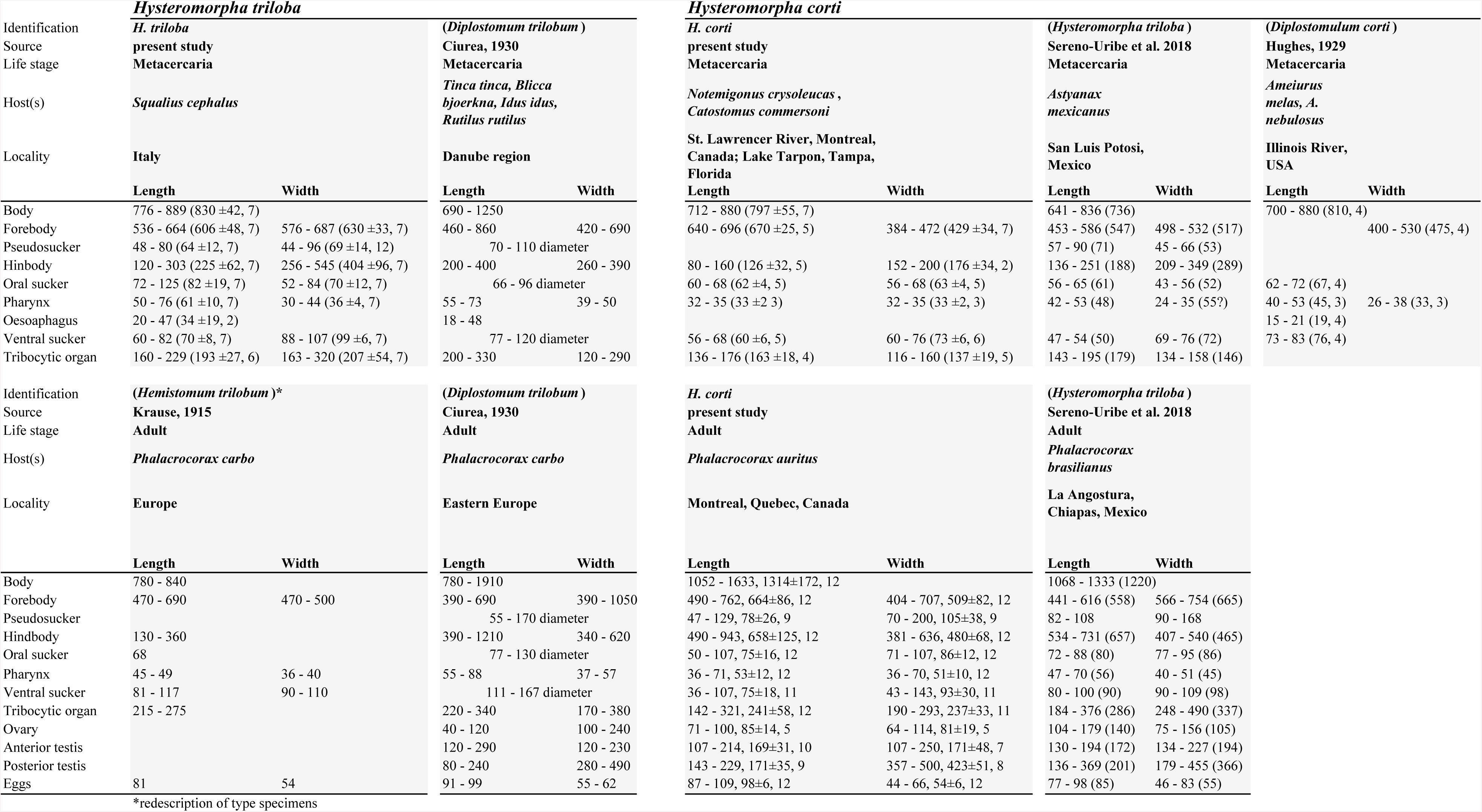
Selected morphometrics from metacercariae and adults of *Hysteromorpha* from the present and other studies, reported in μm as range (mean, ± standard deviation, n).

**Supplementary Table 7:**
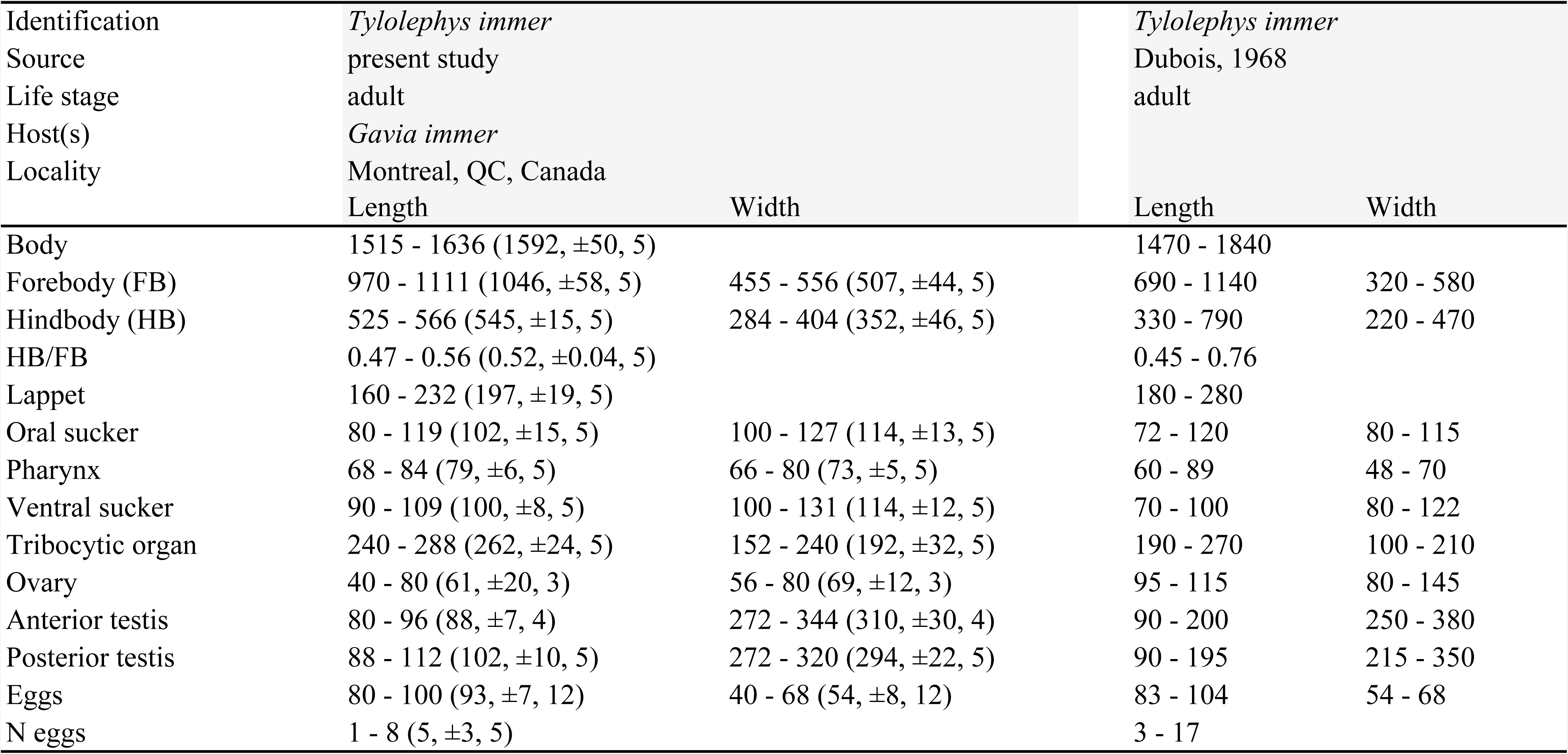
Selected morphometrics from adults of *Tylodelphys immer* reported in μm as range (mean, ± standard deviation, n).

## References

Athokpam, V., Tandon, V., 2014. Morphological and molecular characterization of *Posthodiplostomum* sp. (Digenea: Diplostomidae) metacercaria in the muscles of snakeheads (*Channa punctata*) from Manipur, India. Helminthologia 51, 141–152.

Bernt, M., Donath, A., Jühling, F., Externbrink, F., Florentz, C., Fritzsch, G., Pütz, J., Middendorf, M., Stadler, P.F., 2013. MITOS: improved de novo metazoan mitochondrial genome annotation. Mol. Phylogenet. Evol. 69, 313–319.

Blasco-Costa, I., Faltýnková, A., Georgieva, S., Skírnisson, K., Scholz, T. and Kostadinova, A., 2014. Fish pathogens near the Arctic Circle: molecular, morphological and ecological evidence for unexpected diversity of *Diplostomum* (Digenea: Diplostomidae) in Iceland. Int. J. Parasitol. 44, 703–715.

Blasco-Costa, I., Locke, S.A., 2017. Life history, systematics and evolution of the Diplostomoidea Poirier, 1886: progress, promises and challenges emerging from molecular studies. Adv. Parasitol. 98, 167–244.

Blasco-Costa, I., Poulin, R., Presswell, B., 2016. Morphological description and molecular analyses of *Tylodelphys* sp. (Trematoda: Diplostomidae) newly recorded from the freshwater fish *Gobiomorphus cotidianus* (common bully) in New Zealand. J. Helminthol. 91, 332–345

Blasco-Costa, Isabel, Poulin, R., Presswell, B., 2016. Species of *Apatemon* Szidat, 1928 and *Australapatemon* Sudarikov, 1959 (Trematoda: Strigeidae) from New Zealand: linking and characterising life cycle stages with morphology and molecules. Parasitol. Res. 115, 271–289.

Boone, E.C., Laursen, J.R., Colombo, R.E., Meiners, S.J., Romani, M.F., Keeney, D.B., 2018. Infection patterns and molecular data reveal host and tissue specificity of *Posthodiplostomum* species in centrarchid hosts. Parasitology (In press) DOI:10.1017/S0031182018000306

Brabec, J., Kostadinova, A., Scholz, T., Littlewood, D.T.J., 2015. Complete mitochondrial genomes and nuclear ribosomal RNA operons of two species of *Diplostomum* (Platyhelminthes: Trematoda): a molecular resource for taxonomy and molecular epidemiology of important fish pathogens. Parasit. Vectors 8, 1–11.

Briscoe, A.G., Bray, R.A., Brabec, J., Littlewood, D.T.J., 2016. The mitochondrial genome and ribosomal operon of *Brachycladium goliath* (Digenea: Brachycladiidae) recovered from a stranded minke whale. Parasitol. Int. 65, 271–275.

Brooks, D.R., O’Grady, R.T., Glen, D.R., 1985. Phylogenetic analysis of the Digenea (Platyhelminthes: Cercomeria) with comments on their adaptive radiation. Can. J. Zool. 63, 411–443.

Bunkley-Williams, L., Williams, E.H., 1994. Parasites of Puerto Rican freshwater sport fishes. Department of Natural and Environmental Resources, San Juan, PR.

Bushnell, B., 2014. BBMap: A Fast, Accurate, Splice-Aware Aligner. Presented at the 9th Annual Genomics of Energy & Environment Meeting, USDOE Office of Science (SC), Walnut Creek, CA.

Bykhovskaya-Pavlovskaya, I.E., 1962. Key to Parasites of Freshwater Fish of the U.S.S.R., Keys to Fauna of the U.S.S.R. Izdatelstvo Akademii Nauk SSSR.

Caffara, M., Locke, S.A., Gustinelli, A., Marcogliese, D.J., Fioravanti, M.L., 2011. Morphological and molecular differentiation of *Clinostomum complanatum and *Clinostomum marginatum* (Digenea: Clinostomidae) metacercariae and adults*. J. Parasitol. 97, 884–891.

Campbell, R., 1971. Biology and development of Cotylurus flabelliformis (Trematoda: Strigeidae). PhD Thesis. Iowa State University, Ames, Iowa.

Chaudhary, A., Gupta, S., Verma, C., Tripathi, R., Singh, H.S., 2017. Morphological and molecular characterization of metacercaria of *Tylodelphys* (Digenea: Diplostomidae) from the piscine host, Mystus tengara from India. J. Parasitol. 103, 565–573.

Chen, L., Feng, Y., Chen, H.-M., Wang, L.-X., Feng, H.-L., Yang, X., Mughal, M.-N., Fang, R., 2016. Complete mitochondrial genome analysis of *Clinostomum complanatum* and its comparison with selected digeneans. Parasitol. Res. 115, 3249–3256.

Chibwana, F.D., Blasco-Costa, I., Georgieva, S. Hosead, K.M., Nkwengulila, G., Scholz, T., Kostadinova, A. 2013. A first insight into the barcodes for African diplostomids (Digenea: Diplostomidae): Brain parasites in *Clarias gariepinus* (Siluriformes: Clariidae). Infection, Genetics and Evolution. 17, 62–70.

Ciurea, I., 1930. Contributions à l’étude morphologique et biologique de quelques strigeidés des oiseaux ichtyophages de la faune de Roumanie (recherches expérimentales). Arch. Roum. Pathol. Expérimentale Microbiol. 3, 277–323.

Cort, W.W., 1917. Homologies of the excretory system of the forked-tailed cercariae: A preliminary report. J. Parasitol. 4, 49.

Dubois, G., 1970a. Les fondements de la taxonomie des Strigeata La Rue (Trematoda: Strigeida). Rev. Suisse Zool. 77.

Dubois, G., 1970b. Synopsis des Strigeidae et des Diplostomatidae (Trematoda). Mém. Société Neuchâtel. Sci. Nat. 10, 1–727.

Dubois, G., 1958. Les Strigeida (Trematoda) de Californie de la collection June Mahon. Bull. Société Neuchâtel. Sci. Nat. 81, 69–78.

Dubois, G., 1938. Monographie des Strigeida (Trematoda). Mém. Société Neuchâtel. Sci. Nat. 6, 1–535.

Dzikowski, R., Levy, M.G., Poore, M.F., Flowers, J.R., Paperna, I., 2004. Use of rDNA polymorphism for identification of Heterophyidae infecting freshwater fishes. Dis. Aquat. Organ. 59, 35–41.

Faircloth, B.C., 2016. PHYLUCE is a software package for the analysis of conserved genomic loci. Bioinformatics 32, 786–788.

Faircloth, B.C., Sorenson, L., Santini, F., Alfaro, M.E., 2013. A phylogenomic perspective on the radiation of ray-finned fishes based upon targeted sequencing of Ultraconserved Elements (UCEs). PLoS ONE 8, e65923.

Fraija-Fernández, N., Olson, P.D., Crespo, E.A., Raga, J.A., Aznar, F.J., Fernández, M., 2015. Independent host switching events by digenean parasites of cetaceans inferred from ribosomal DNA. Int. J. Parasitol. 45, 167–173.

Galazzo, D.E., Dayanandan, S., Marcogliese, D.J., McLaughlin, J.D., 2002. Molecular systematics of some North American species of *Diplostomum* (Digenea) based on rDNA-sequence data and comparisons with European congeners. Can. J. Zool. 80, 2207–2217.

Gibson, D.I., 1996. Guide to the Parasites of Fishes of Canada: Trematoda, Canadian Special Publication of Fisheries and Aquatic Sciences. NRC Research Press.

Gibson, D.I., 1987. Questions in digenean systematics and evolution. Parasitology 95, 429–460.

Gibson, D.I., Bray, R.A., 1994. The evolutionary expansion and host-parasite relationships of the Digenea. Int. J. Parasitol. 24, 1213–1226.

Graybeal, A., Cannatella, D., 1998. Is it better to add taxa or characters to a difficult phylogenetic problem? Syst. Biol. 47, 9–17.

Hatch, J.J., Brown, K.M., Hogan, G.G., Morris, R.D., 2000. Great Cormorant (Phalacrocorax carbo). Birds N. Am. Online.

Hedtke, S.M., Townsend, T.M., Hillis, D.M., Collins, T., 2006. Resolution of phylogenetic conflict in large data sets by increased taxon sampling. Syst. Biol. 55, 522–529.

Hernández-Mena, D.I., García-Prieto, L., García-Varela, M., 2014. Morphological and molecular differentiation of *Parastrigea* (Trematoda: Strigeidae) from Mexico, with the description of a new species. Parasitol. Int. 63, 315–323.

Hernández-Mena, D.I., García-Varela, M., Pérez-Ponce de León, G., 2017. Filling the gaps in the classification of the Digenea Carus, 1863: systematic position of the Proterodiplostomidae Dubois, 1936 within the superfamily Diplostomoidea Poirier, 1886, inferred from nuclear and mitochondrial DNA sequences. Syst. Parasitol. 94, 833–848.

Hoffman, G.L., 1958. Experimental studies on the cercaria and metacercaria of a strigeoid trematode, Posthodiplostomum minimum. Exp. Parasitol. 7, 23–50.

Howe, K.L., Bolt, B.J., Cain, S., Chan, J., Chen, W.J., Davis, P., Done, J., Down, T., Gao, S., Grove, C., Harris, T.W., Kishore, R., Lee, R., Lomax, J., Li, Y., Muller, H.-M., Nakamura, C., Nuin, P., Paulini, M., Raciti, D., Schindelman, G., Stanley, E., Tuli, M.A., Van Auken, K., Wang, D., Wang, X., Williams, G., Wright, A., Yook, K., Berriman, M., Kersey, P., Schedl, T., Stein, L., Sternberg, P.W., 2016. WormBase 2016: expanding to enable helminth genomic research. Nucleic Acids Res. 44, D774–D780.

Hugghins, E.J., 1954a. Life history of a strigeid trematode, *Hysteromorpha triloba* (Rudolphi, 1819) Lutz, 1931. II. Sporocyst through adult. Trans. Am. Microsc. Soc. 73, 221–236.

Hugghins, E.J., 1954b. Life history of a strigeid trematode, Hysteromorpha triloba Rudolphi, 1819) Lutz, 1931. I. Egg and miracidium. Trans. Am. Microsc. Soc. 73, 1–15.

Hughes, R.C., 1929. Studies on the trematode family Strigeidae (Holostomidae), No. XIV: Two new species of Diplostomula. Occas. Pap. Mus. Zool. 202, 1–29.

Inoue, J.G., Miya, M., Tsukamoto, K., Nishida, M., 2003. Basal actinopterygian relationships: a mitogenomic perspective on the phylogeny of the “ancient fish.” Mol. Phylogenet. Evol. 26, 110–120.

Johnson, A.D., 1970. Alaria mustelae: Description of mesocercaria and key to related species. Trans. Am. Microsc. Soc. 89, 250–253.

Kalbe, M., Wegner, K.M., Reusch, T.B.H., 2016. Dispersion patterns of parasites in 0+ year three-spined sticklebacks: a cross population comparison. J. Fish Biol. 60, 1529–1542.

Kostadinova, A., Pérez-del-Olmo, A., 2014. The Systematics of the Trematoda, in: Toledo, R., Fried, B. (Eds.), Digenetic Trematodes. Springer New York, New York, NY, pp. 21–44.

Kvach, Y., Jurajda, P., Bryjová, A., Trichkova, T., Ribeiro, F., Přikrylová, I., Ondračková, M., 2017. European distribution for metacercariae of the North American digenean *Posthodiplostomum* cf. minimum centrarchi (Strigeiformes: Diplostomidae). Parasitol. Int. 66, 635–642.

Kvach, Y., Ondračková, M., Jurajda, P., 2016. First report of metacercariae of *Cyathocotyle prussica* parasitising a fish host in the Czech Republic, Central Europe. Helminthologia 53, 1–5.

La Rue, G.R., 1957. The classification of Digenetic Trematoda: A review and a new system. Exp. Parasitol. 6, 306–349.

Lapage, G., 1961. A list of the parasitic protozoa, helminths and Arthropoda recorded from species of the family Anatidae (ducks, geese and swans). Parasitology 51, 1–109.

Locke, S.A., Al-Nasiri, F.S., Caffara, M., Drago, F., Kalbe, M., Lapierre, A.R., McLaughlin, J.D., Nie, P., Overstreet, R.M., Souza, G.T., Takemoto, R.M., Marcogliese, D.J., 2015. Diversity, specificity and speciation in larval Diplostomidae (Platyhelminthes: Digenea) in the eyes of freshwater fish, as revealed by DNA barcodes. Int. J. Parasitol. 45, 841–855.

Locke, S.A., McLaughlin, J.D., Lapierre, A.R., Johnson, P.T., Marcogliese, D.J., 2011. Linking larvae and adults of *Apharyngostrigea cornu, Hysteromorpha triloba* and *Alaria mustelae* (Diplostomoidea: Digenea) using molecular data. J. Parasitol. 97, 846–851.

Locke, S.A., McLaughlin, J.D., Marcogliese, D.J., 2010. DNA barcodes show cryptic diversity and a potential physiological basis for host specificity among Diplostomoidea (Platyhelminthes: Digenea) parasitizing freshwater fishes in the St. Lawrence River, Canada. Mol. Ecol. 19, 2813–2827.

López-Hernández, D., Locke, S.A., de Melo, A.L., Leite Rabelo, E., Pinto, H.A., 2018. Molecular, morphological and experimental assessment of the life cycle of *Posthodiplostomum nanum* Dubois 1937 (Trematoda: Diplostomidae) from Brazil, with phylogenetic evidence of paraphyly of the genus *Posthodiplostomum* Dubois, 1936. Infect. Genet. Evol. In press. DOI:10.1016/j.meegid.2018.05.010

López-Jiménez, A., Pérez-Ponce de León, G., García-Varela, M., 2017. Molecular data reveal high diversity of *Uvulifer* (Trematoda: Diplostomidae) in Middle America, with the description of a new species. J. Helminthol. In press. DOI:10.1017/S0022149X17000888

Lunaschi, L.I., Cremonte, F., Drago, F.B., 2007. Checklist of digenean parasites of birds from Argentina. Zootaxa 1403, 1–36.

Lunter, G., Goodson, M., 2011. Stampy: a statistical algorithm for sensitive and fast mapping of Illumina sequence reads. Genome Res. 21, 936–939.

Lutz, A., 1931. Contribuição ao conhecimento da ontogenia das strigeidas. Mem. Inst. Oswaldo Cruz 25, 333–342.

MacCallum, G., 1921. Studies in helminthology. Zoopathologica 1, 137–284.

McDonald, M.E., 1981. Key to Trematodes Reported in Waterfowl. US Fish and Wildlife Service.

Moszczynska, A., Locke, S.A., McLaughlin, J.D., Marcogliese, D.J., Crease, T.J., 2009. Development of primers for the mitochondrial cytochrome *c* oxidase I gene in digenetic trematodes (Platyhelminthes) illustrates the challenge of barcoding parasitic helminths. Mol. Ecol. Resour. 9, 75–82.

Niewiadomska, K., 2002a. Superfamily Diplostomoidea, in: Gibson, D.I., Jones, A., Bray, R. (Eds.), Keys to the Trematoda. CABI Publishing, Oxon, UK, pp. 159–166.

Niewiadomska, K., 2002b. Family Strigeidae Railliet, 1919, in: Gibson, D., Jones, A., Bray, R. (Eds.), Keys to the Trematoda. CABI Publishing, Oxon, UK, pp. 231–241.

Niewiadomska, K., 2002c. Family Diplostomidae Poirier, 1886, in: Gibson, D.I., Jones, A., Bray, R. (Eds.), Keys to the Trematoda. CABI Publishing, Oxon, UK, pp. 167–196.

Olson, P., Cribb, T., Tkach, V., Bray, R., Littlewood, D., 2003. Phylogeny and classification of the Digenea (Platyhelminthes: Trematoda). Int. J. Parasitol. 33, 733–755.

Overstreet, R.M., Curran, S.S., Pote, L.M., King, D.T., Blend, C.K., Grater, W.D., 2002. Bolbophorus damnificus n. sp. (Digenea: Bolbophoridae) from the channel catfish *Ictalurus punctatus* and American white pelican *Pelecanus erythrorhynchos* in the USA based on life-cycle and molecular data. Syst. Parasitol. 52, 81–96.

Palmieri, J.R., 1977. Host-induced morphological variations in the strigeoid trematode *Posthodiplostomum minimum* (Trematoda: Diplostomatidae) II: Body measurements and tegument modifications. Gt. Basin Nat. 37, 129–137.

Park, G.-M., 2007. Genetic comparison of liver flukes, *Clonorchis sinensis* and *Opisthorchis viverrini*, based on rDNA and mtDNA gene sequences. Parasitol. Res. 100, 351–357.

Pearson, J.C., 1972. A phylogeny of life-cycle patterns of the Digenea. Adv. Parasitol. 10, 153–189.

Pearson, J.C., Johnson, A.D., 1988. The taxonomic status of *Alaria marcianae* (Trematoda: Diplostomidae). Proc. Helminthol. Soc. Wash. 55, 102–103.

Peng, Y., Leung, H.C., Yiu, S.-M., Chin, F.Y., 2012. IDBA-UD: a de novo assembler for single-cell and metagenomic sequencing data with highly uneven depth. Bioinformatics 28, 1420–1428.

Perea, S., Vukić, J., Šanda, R., Doadrio, I., 2016. Ancient mitochondrial capture as factor promoting mitonuclear discordance in freshwater fishes: A case study in the genus *Squalius* (Actinopterygii, Cyprinidae) in Greece. PLOS ONE 11, e0166292.

Philippe, H., Brinkmann, H., Lavrov, D.V., Littlewood, D.T.J., Manuel, M., Wörheide, G., Baurain, D., 2011. Resolving difficult phylogenetic questions: Why more sequences are not enough. PLoS Biol. 9, e1000602.

Platt, R.N., Faircloth, B.C., Sullivan, K.A., Kieran, T.J., Glenn, T.C., Vandewege, M.W., Lee, T.E., Baker, R.J., Stevens, R.D., Ray, D.A., 2018. Conflicting evolutionary histories of the mitochondrial and nuclear genomes in New World *Myotis* bats. Syst. Biol. 67, 236–249.

Pleijel, F., Jondelius, U., Norlinder, E., Nygren, A., Oxelman, B., Schander, C., Sundberg, P., Thollesson, M., 2008. Phylogenies without roots? A plea for the use of vouchers in molecular phylogenetic studies. Mol. Phylogenet. Evol. 48, 369–371.

Rudolphi, C., 1819. Entozoorum synopsis cui accedunt mantissima duplex et indices locupletissima. Sumptibus Augusti Rücker, Berlin, Germany

Sattmann, H., Hörweg, C., Stagl, V., 2014. Johann Gottfried Bremser (1767–1827) und die Kuhpockenimpfung. Wien. Klin. Wochenschr. 126, 3–10.

Sereno-Uribe, A.L., López-Jimenez, A., Andrade-Gómez, L., García-Varela, M., 2018. A morphological and molecular study of adults and metacercariae of *Hysteromorpha triloba* (Rudolpi, 1819), Lutz 1931 (Diplostomidae) from the Neotropical region. J. Helminthol. In press. DOI:10.1017/S0022149X17001237

Shoop, W.L., 1989. Systematic analysis of the Diplostomidae and Strigeidae (Trematoda). J. Parasitol. 75, 21–32.

Silvestro, D., Michalak, I., 2012. raxmlGUI: a graphical front-end for RAxML. Org. Divers. Evol. 12, 335–337.

Simão, F.A., Waterhouse, R.M., Ioannidis, P., Kriventseva, E.V., Zdobnov, E.M., 2015. BUSCO: assessing genome assembly and annotation completeness with single-copy orthologs. Bioinformatics btv351.

Stamatakis, A., 2014. RAxML version 8: a tool for phylogenetic analysis and post–analysis of large phylogenies. Bioinformatics 30, 1312–1313.

Stoyanov, B., Georgieva, S., Pankov, P., Kudlai, O., Kostadinova, A., Georgiev, B.B., 2017. Morphology and molecules reveal the alien *Posthodiplostomum centrarchi* Hoffman, 1958 as the third species of *Posthodiplostomum* Dubois, 1936 (Digenea: Diplostomidae) in Europe. Syst. Parasitol. 94, 1–20.

Stunkard, H.W., 1946. Interrelationships and taxonomy of the digenetic trematodes. Biol. Rev. 21, 148–158.

Sun, M., Soltis, D.E., Soltis, P.S., Zhu, X., Burleigh, J.G., Chen, Z., 2015. Deep phylogenetic incongruence in the angiosperm clade Rosidae. Mol. Phylogenet. Evol. 83, 156–166.

Tamura, K., Stecher, G., Peterson, D., Filipski, A., Kumar, S., 2013. MEGA6: Molecular Evolutionary Genetics Analysis version 6.0. Mol. Biol. Evol. 30, 2725–2729.

Uhrig, E.J., Spagnoli, S.T., Tkach, V.V., Kent, M.L., Mason, R.T., 2015. Alaria mesocercariae in the tails of red-sided garter snakes: evidence for parasite-mediated caudectomy. Parasitol. Res. 114, 4451–4461.

Waeschenbach, A., Webster, B.L., Littlewood, D.T.J., 2012. Adding resolution to ordinal level relationships of tapeworms (Platyhelminthes: Cestoda) with large fragments of mtDNA. Mol. Phylogenet. Evol. 63, 834–847.

Whitfield, J.B., Lockhart, P.J., 2007. Deciphering ancient rapid radiations. Trends Ecol. Evol. 22, 258–265.

Zeder, J., 1800. Erster Nachtrag zur Naturgeschichte der Eingeweidewürmer. Leipzig.

